# Altered Mechanobiology of PDAC Cells with Acquired Chemoresistance to Gemcitabine and Paclitaxel

**DOI:** 10.1101/2024.04.10.588671

**Authors:** Alessandro Gregori, Cecilia Bergonzini, Mjriam Capula, Rick Rodrigues de Mercado, Erik H. J. Danen, Elisa Giovannetti, Thomas Schmidt

## Abstract

**Background:** Pancreatic ductal adenocarcinoma (PDAC) acquired resistance to chemotherapy poses a major limitation to patient survival. Despite understanding of some biological mechanisms of chemoresistance, much of those mechanisms remain to be uncovered. Mechanobiology, which studies physical properties of cells, holds promise as a potential target for addressing challenges of chemoresistance in PDAC. Therefore, we here in an initial step, assessed the altered mechanobiology of PDAC cells with acquired chemoresistance to gemcitabine and paclitaxel.

**Methods:** Five PDAC cell lines and six stably-resistant subclones were assessed for force generation on elastic micropillar arrays. Those measurements of mechanical phenotype were complemented by single-cell motility and invasion in collagen matrix were investigated using 2D models and 3D extracellular matrix-mimetic, respectively. Further the nuclear translocation of Yes-associted protein (YAP), as a measure of active mechanical status, was compared, and biomarkers of the epithelial-to-mesenchymal transition (EMT) were evaluated using RT-PCR.

**Results:** PDAC cells with acquired chemoresistance exert higher traction forces than their parental/wild-type (WT) cells. In 2D, single-cell motility was altered for all chemoresistant cells, with a cell-type specific pattern. In 3D, spheroids of chemoresistant PDAC cells were able to invade the matrix, and remodel collagen more than their WT clones. However, YAP nuclear translocation and EMT were not significantly altered in relation to changes in other physical parameters.

**Conclusion:** This is the first study to investigate and report on the altered mechanobiological features for PDAC cells that have acquired chemoresistance. A better understanding of mechanical features could help in identifying future targets to overcome chemoresistance in PDAC.

## Introduction

Pancreatic ductal adenocarcinoma (PDAC) is the most common type of pancreatic cancer, with a dismal prognosis. The 5-year survival is reached by only 13% of all patients (see 2024 update at: Pancreatic Cancer Five-Year Survival Rate Increases to 13% - Pancreatic Cancer Action Network (pancan.org)) [1]. The lack of early diagnosis, limited therapeutic options and inherent or acquired chemoresistance all concur to the poor prognosis [2]. Besides surgical resection, for which a small portion of patients are eligible, chemotherapy is the only available treatment. Currently, the combination-regimens FOLFIRINOX or gemcitabine plus nab-paclitaxel are being employed to treat patients with inoperable PDAC [3]. However, in the majority of the patients rapid chemoresistance occurs, with the underlying biological mechanisms remaining largely elusive. Therefore, mechanobiology, which analyzes physical forces in a biological context, is gaining attention in the studies on chemoresistance of PDAC [4].

Cellular mechano-transduction, the process by which mechanical cues are converted into intracellular molecular signals, along with interactions with the surrounding extracellular matrix (ECM) play a pivotal role in cell homeostasis and signaling. Increasing evidence show that this phenomenon is particularly critical in cancers that develop abnormal ECM, causing tumor progression, increased metastatic potential and drug resistance [5–7]. PDAC is characterized by a drastic change in the ECM reflected by tissue stiffening. Indeed, PDAC progression is accompanied by a desmoplastic reaction, which produces a dense stroma constituting up to 90% of the tumor volume, and increases tissue stiffness up to 50 kPa in terms of Young’s modulus [8]. Therefore, PDAC cells are subject to many mechanical stimuli which are thought to promote tumor progression and therapy resistance. One such mechanism that we previously reported is overexpression of the integrin alpha 2 (ITGA2), a cellular stiffness sensor, that promotes gemcitabine resistance in PDAC [9]. Additionally, Rice and collaborators showed involvement of epithelial-to-mesenchymal transition (EMT) in inducing chemoresistance of PDAC cells [10].

Metastatic dissemination and cell migration are widely affected by stromal stiffness in many cancers [11]. In general, cells generate forces to migrate through the ECM and a correlation between metastatic potential and increased force generation has been previously described for many types of cancers [12]. Likewise, it is believed that EMT is required for cells to loose ECM adhesion and migrate [13]. EMT has been investigated in PDAC in relation to metastasis and inherent or acquired chemoresistance [10,13–15]. Induction of EMT, and activation of cellular forces are translated into mechanical stimuli, that in turn trigger Yes-associated protein (YAP) nuclear translocation and downstream signaling [10].

Despite those general observations, there is limited understanding regarding whether cells with acquired chemoresistance exhibit a deregulated mechano-transduction. We hypothesized that exposing PDAC cells to paclitaxel, a drug that inhibits microtubule disassembly, impacts cytoskeleton activity and/or mechanical signaling. Consequently, we here investigated and report, for the first time, that PDAC cells with acquired chemoresistance to gemcitabine and paclitaxel display an altered mechanical phenotype. For investigation, the present study assessed force generation in conditions of varying stiffness in 5 PDAC cell lines, and in 6 chemoresistant subclones. Additionally, singe-cell motility in 2D and 3D collagen-embedded spheroids were investigated to determine the invasive potential. Finally, Hippo signaling and EMT-mediators of mechano-transduction were analyzed, revealing that mechanical forces in chemoresistant cells were accompanied by YAP nuclear translocation, and changes in the expression of EMT genes.

## Methods

### Cell culture

One non-tumor cell line, HPDE kindly provided by Ming Tsao (Ontario Cancer Institute, Toronto, Ontario, Canada), and five PDAC cell lines were employed in this study: CAPAN-1 (epithelial phenotype) and BxPC-3 (epithelial phenotype) were purchased from ATCC, PATU-T (mesenchymal phenotype) were kindly provided by Dr. Irma van Die (Amsterdam UMC, Amsterdam, The Netherland), SUIT-2.028 and SUIT-2.007 (epithelial and mesenchymal phenotype, respectively) were kindly provided by Dr. Adam Frampton (Imperial College London, London, UK). All gemcitabine-resistant and paclitaxel-resistant cells were generated by continuous drug exposure over a period of 12 months, as previously described [16]. HPDE were cultured in KGM medium, PATU-T in DMEM and all other cell lines in RPMI medium. All cells were supplemented with 10% heat-inactivated new-born calf serum and 1% penicillin/streptomycin and kept at 37 °C and 5% CO_2_. Cells were periodically tested for mycoplasma contamination.

### Elastic Micropillar arrays

In this study, we used elastic micropillar arrays (μPA) of polydimethylsiloxane (PDMS, Sylgard 184, Dow Corning) arranged in hexagonal geometry, of 2 μm diameter, 4 μm spacing, and height (effective Young’s modulus) of 3.2 μm (11 kPa), 4.1 μm (29 kPa), 6.1 μm (47 kPa) and 6.9 μm (142 kPa), respectively. μPAs were generated as previously described in detail [17,18]. Briefly, a 1:10 PDMS mixture (crosslinker/base ratio) was poured into a negative mold made in silicon-wafers, and cured for 20 hours at 110 °C. Next, μPA were peeled off the wafers, activated with ultraviolet light for 10 minutes and coated with fibronectin (F1141; Sigma-Aldrich) using micro-contact printing. A mixture of 1:5 AlexaFluor488-labeled to unlabeled fibronectin was used for coating the top of the pillar arrays.

### Cell seeding, immunostaining and microscopy

Cells were seeded on μPAs and incubated at 37 °C for 16-19 hours, in order to allow for attachment and spreading, but before duplication. Subsequently, cells were fixed for 15 minutes in 4% paraformaldehyde, permeabilized 10 minutes in 0.1% Triton X-100, blocked for 1 hour in 5% BSA and stained for 1 hour with AlexaFluor561-Phalloidin. Finally, nuclei were counterstained with DAPI. μPAs were then flipped upside-down on a 25 mm diameter #1.5 coverglass, and imaged using a 100x oil-immersion objective on an Axiovert200 optical microscope (Carl Zeiss, Oberkochen, Germany) equipped with a spinning disk unit (CSU-X1; Yokogawa Electric, Musashino, Tokyo, Japan), and an emCCD camera (iXon 897; Andor Labs, Morrisville, NC, USA).

### Pillars deflection analysis

Pillar deflection was analyzed using a custom-designed Matlab (Matlab R2018a; MathWorks, Natick, MA, USA) script, as previously described [17]. Briefly, the center of each pillar was determined to a precision of ∼50 nm. Pillar displacements from the original positions within a hexagonal grid were used to map the force-induced pillar-deflections (Δx). The force was calculated from the deflection and the pillars elastic constant (k) using Hooke’s law: *F* = *k* * Δ*x*.

Traction forces were defined as the inward-pointing forces (F_*in*_). All outward-pointing forces were excluded from further analyses.

### YAP nuclear translocation

Cells were seeded on μPA and immunostained as described above. Cells were incubated for 1 hour with a Yes-associated protein (YAP) primary antibody (Santa Cruz Biotechnology, sc101199, 1:200) followed by secondary antibody labeling. Cells were then imaged on a Nikon TEi2 confocal microscope equipped with automated stage, and controlled by NIS Element software (Nikon). YAP nuclear translocation was calculated as the percentage of nuclear YAP over the total YAP:

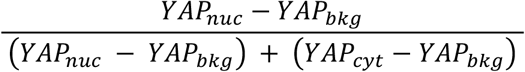

Where YAP_nuc_ is nuclear YAP, YAP_bkg_ is the background in the YAP channel outside the cell area, and YAP_cyt_ is cytosolic YAP. Correlation between YAP and mean force per pillar was calculated in R Studio using the linear regression model of the *ggplot* package.

### Quantitative PCR (RT-qPCR)

Cells were seeded in 6-well plates (VWR). RNA was extracted from adherent cells 48 hours post-seeding using TriZol Reagent (Invitrogen) according to the manufacturer’s instructions. 500 ng of RNA was used to retrotranscribe cDNA with the First Strand cDNA synthesis kit (ThermoScientific, #K1612), following manufacturer instructions. RT-qPCR was carried out with Sso Advanced Universal SYBR Green Supermix (Bio-Rad, #172-5271) in a CFX96 Real-Time System (Bio-Rad) apparatus. Primer sequences (Invitrogen) for E-cadherin (*E-cad*), N-cadherin (*N-cad*), Vimentin (*Vim*) and Beta-actin (*β-actin*) can be found in the Supplemental Table 1.

### Single cell motility

96-well Screenstar black µClear plates (#655866) were coated with 20 µg/µL rat-tail collagen (Ibidi) or 20 µg/µL human fibronectin in Ultrapure water for 1 hour at 37 °C. After removing the coating solution, coated wells were washed three times with PBS. PDAC cells were seeded at different densities: PATU-T (6000 cells/well); SUIT-2.007 and SUIT-2.028 (7000 cells/well). After 24 hours, all cell lines were treated with 5 µM Verapamil or medium (control) for 1 hour. Verapamil was used to allow Hoechst accumulation in PR cells, which overexpress ABCB1, as previously described [16]. Cells were incubated with Hoechst 33342 solution for 1 hour, then medium was refreshed. A concentration of 1.25 µg/µL bosutinib was used as positive control for inhibited migration and added to selected wells before imaging. Plates were then imaged with a 20x objective on a ImageXpress micro XLS imager (Molecular Devices), equipped with an incubator to maintain cells at 37 °C and 5% CO_2_. Four fields per well were imaged, every 12 minutes, for 16 hours. Two wells per conditions were analyzed, except for bosutinib which was tested in single replicate. Data from at least three experiments were collected. Images were analyzed with a Matlab script as previously described [19], after adaptation to identify and localize cell nuclei. Cells were identified and localized from thresholded images. Noise (object size <1 μm^2^) and large clusters (object size >9 μm^2^) were discarded from further analysis. From the position data, trajectories were determined and mobility analysis applied. Cell mobility was analyzed in terms of the change in their mean-squared displacement (*msd*) with lag time (*t*_*lag*_) between two observations. For the analysis we assumed both a diffusive motion characterized by a diffusion constant (*D*), and directed motion characterized by a velocity (*v*) (for further details consider the supplemental material). Subsequently, the diffusive fraction of the motility pattern was characterized by a diffusive fraction (*f*_*D*_) which we defined as the ratio of the diffusive part of the *msd* to the total *msd*, at fixed lagtime time, *t*_*D*_ = 10 s.

Each trajectory was characterized by those three parameter. From the single-trajectory analysis means and standard deviations of the cell population were subsequently determined as presented in the results section. The equations describing mean-squared displacement and diffusive fraction can be found in the supplementary material.

### 3D ECM remodeling ability

PDAC spheroids embedded in 1 mg/mL rat-tail type-I collagen gels were obtained by automated microinjection as previously described [9,20]. The gels were polymerized in 384-well µclear plates (Greiner) at 37 °C for 1 hour, washed with growth medium after polymerization for 6 times every 15 minutes, and 1 time for 1 hour, before cell injection. Images of tumor spheroids were acquired using a Nikon Eclipse TEi2 inverted scanning confocal microscope equipped with laser lines 405 nm, 488 nm, 561 nm and 640 nm, an A1R MP scanner, a Nikon encoded and automated stage and temperature and CO_2_-controlled incubator. The microscope was controlled through NIS Element Software (Nikon Instruments Inc., Melville, NY, USA). Images were acquired after 24 hours with a Plan Apo 20×/0.75 NA objective (Nikon Instruments Inc., Melville, NY, USA). For reflection microscopy, collagen fibers were scanned at 561 nm excitation with a 561-blocking dichroic mirror. All other wavelengths passed a bandpass filter 400-750 nm. The image-stitching function from the software was used to combine 2 x 2 images. At least 19 z-planes, 5 µm apart were acquired for each spheroid. To analyze the invasion/migration potential of PDAC spheroids samples were fixed, permeabilized and stained 48 hours post injection with a solution containing: 0.05 µM rhodamine phalloidin (Invitrogen), 2 µg/mL Hoechst33342, Triton-X 0.1% (Sigma), 2% PFA and PBS. After an O/N incubation at 4 °C, gels were washed 3 times with PBS, and images of the spheroids were captured with a Nikon TEi2 confocal microscope, and a 20x objective. To obtain the migration area, the z-projection of the spheroid images were analyzed using a Matlab script described previously. Briefly, z-stack images of spheroids were projected using the standard deviation in the z-direction of the rhodamine phalloidin channel. The foreground was separated from the background using an adaptive threshold. Morphological features, such as the area, were extracted from the projected foreground of the individual spheroids. Images of day 0 were acquired with a phase-contrast microscope connected to a camera, and the area was measured with ImageJ after conversion to 8-bit, and thresholding to select only the area of the spheroid. Both day-2 and day-0 area values were converted to µm^2^ in order to calculate the relative area as the ratio between day-2 and day-0. Alignment of collagen fibers was calculated using the CurveAlign software [22]. First, the CT-FIRE module was used to identify collagen fibers in each z-plane. Then, CurveAlign was used in the CT-Fire Segment mode, to remove noise from the image and enhance the fiber edges through the curvelet transform, and then identify the fiber network through a fiber tracking algorithm. Subsequently, fibers orientation with respect to the spheroid edge are measured (boundary analysis). In particular, the mask of the core of the spheroid is provided to the software, which then calculates the relative angle of the fibers with the tangent to the closest point of the boundary. The relative angles are categorized in sectors of 5 degrees, from 2.5 to 87.5, with 2.5 being almost parallel to the tangent to the boundary, and 87.5 being perpendicular to it. Finally, output data from all the z-planes of at least 4 spheroids, in 2 biological replicates, were combined and analyzed with R Studio to create polar plots, showing the distribution of the % of the relative angles for each cell line. For each spheroid, the number of fibers for each degree range was summed through all the z-planes of one spheroid. Then, the % of fibers in each degree range over the total number of fibers was calculated. GraphPad Prism version 9 (Intuitive Software for Science, San Diego, CA, USA) was used for displaying the % of aligned fibers per spheroid, which we defined as the % of fibers with relative angles between 72.5 and 90 degrees.

### Statistical analysis

All pillar experiments were performed at least in biological triplicate with more than 20 cells analyzed per experimental run. Wilcoxon test was performed in R Studio to compare cell spreading area and mean force per pillar among the different conditions; linear regression was assessed in R Studio. Statistical significance of % of nuclear YAP and metastatic separation of mean force/pillar were assessed with GraphPad Prism using one-way ANOVA multiple comparison Tukey’s test. Statistical significance of migration velocity, diffusive fraction, diffusion constant, collagen alignment and relative area were assessed with GraphPad Prism with ordinary one-way ANOVA multiple comparison test followed by a Šídák’s post-hoc test.

## Results

In order to elucidate the mechanical properties of PDAC cells we selected a panel of commercially available cell lines, assessing their force generation using elastic micropillar arrays (μPA) of varying stiffness. Subsequently we generated a panel of chemoresistant cells and evaluated whether those have altered mechanical characteristics in terms of force generation, YAP nuclear translocation, EMT, single-cell motility and 3D invasion capacity in collagen (see the analysis pipeline in Figure 1).

**Figure 1.**
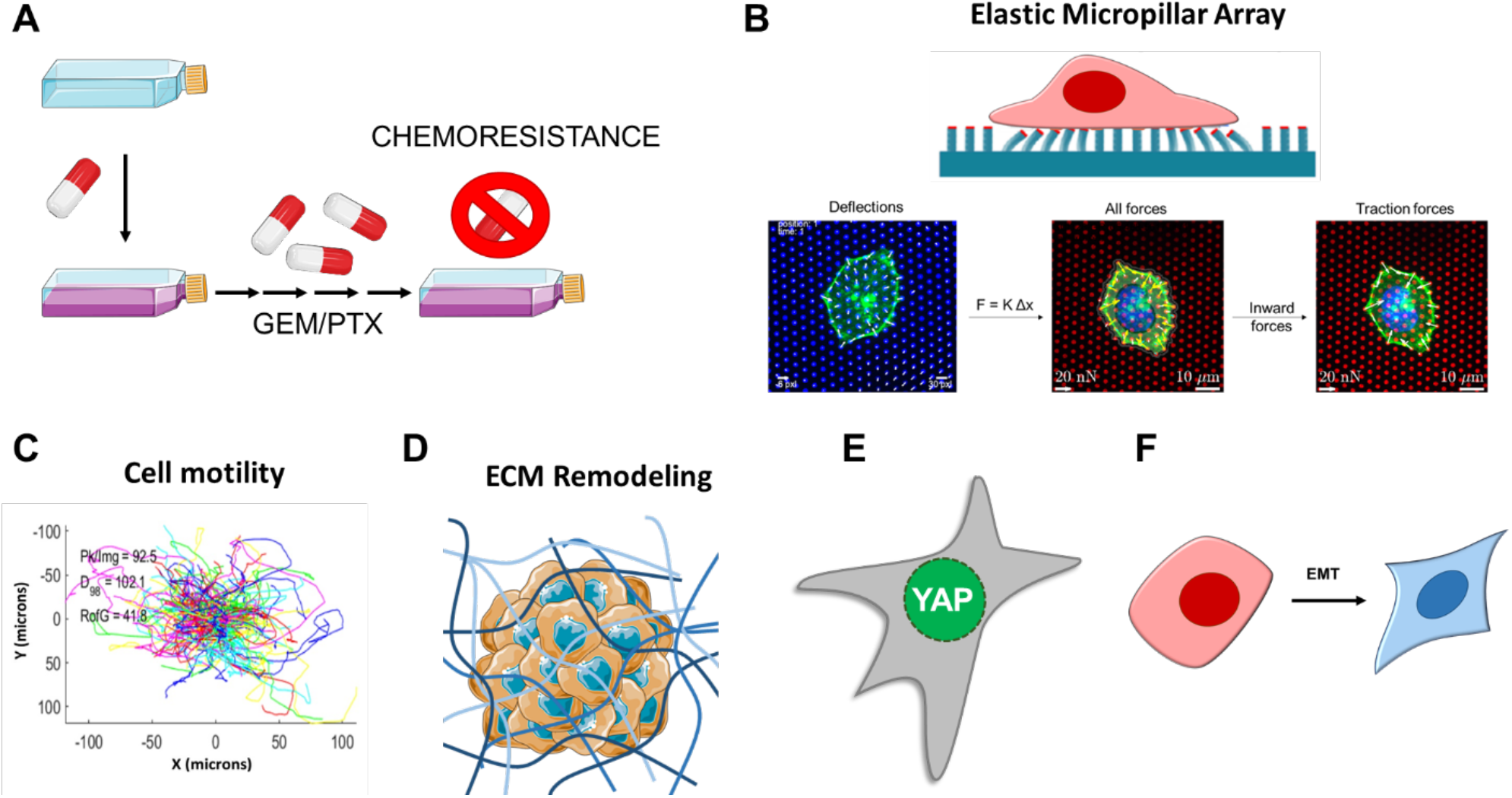
Workflow of the study. (**A**) PDAC cells were exposed to gemcitabine (GEM) or paclitaxel (PTX) to generate chemoresistant clones. (**B**) PDAC cells, and their resistant subclones, seeded on elastic micropillar arrays of varying stiffness were assessed for force generation by measuring the pillar deflections. Traction forces were defined as the inward-pointing forces. (**C**) Single-cell motility was assessed in cells seeded on collagen and fibronectin-coated substrates. (**D**) 3D collagen-embedded spheroid invasion, and spheroid-induced ECM remodeling was analyzed. (**E**) YAP nuclear translocation assessed by immunofluorescence for cells growing on soft pillars. (**F**) Biomarkers of epithelial-to-mesenchymal transition (EMT) were assessed by RT-qPCR. Part of the figure was adapted from images made by Servier Medical Art by Servier, licensed under a Creative Commons Attribution 4.0 Unported License, at https://smart.servier.com.

We selected a heterogeneous panel of commercially available cell lines including: one non-tumor cell line, HPDE (human pancreatic ductal epithelial), three epithelial-phenotype cell lines BxPC-3, SUIT-2.028 and CAPAN-1 (with BxPC-3 being *KRAS* wild-type), and one mesenchymal-phenotype cell SUIT-2.007. μPA of varying stiffness, coated with fibronectin, were employed to assess cellular force generation. The height of the pillars were 10, 8, 6, 4 μm which resulted in a variation of the effective stiffness of the surface of 11, 29, 47 and 142 kPa (Young’s modulus), respectively [17].

### The spreading area of PDAC cells varies among cell lines, and with substrate stiffness

We first evaluated whether the cell spreading area was affected by pillar stiffness, and whether different cell lines had similar spreading area. As has been observed for a variety of cell lines [23,24], we confirmed that also PDAC cells assume a larger spreading area on stiffer plating conditions (Figure 2 and Supplemental Table 2). Additionally, three cell lines, HPDE and SUIT-2.007/028, showed a larger area than CAPAN-1 and BxPC-3 in all stiffness conditions (Figure 2B). PDAC cells that had a larger spreading area consequently deflected more pillars.

**Figure 2.**
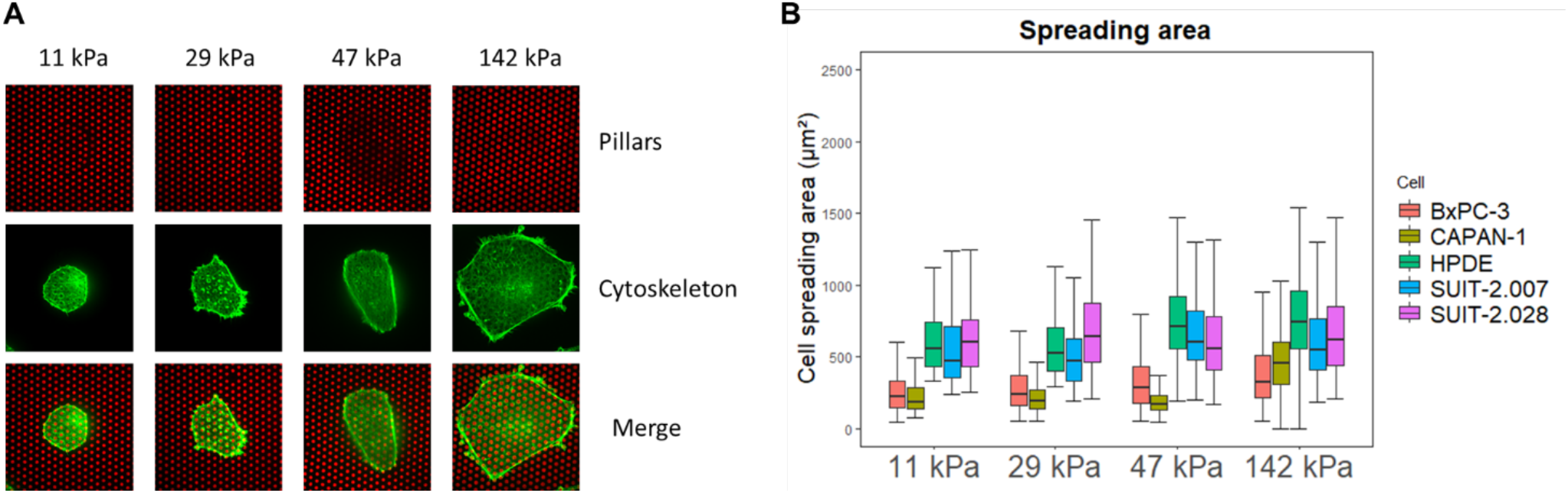
PDAC cell spreading area varies with stiffness. (**A**) Representative confocal microscopy images of BxPC-3 cells growing on fibronectin-coated pillars of different stiffness. (**B**) Boxplots of cell spreading area (μm^2^) for BxPC-3, CAPAN-1, HPDE, SUIT-2.028, and SUIT-2.007 growing on fibronectin-coated pillars (25^th^ and 75^th^ percentiles marked, line at median).

### PDAC cells force generation is stiffness-dependent

The total force applied by one cell is determined by the sum of all the forces on all deflected pillars beneath the cell spreading area. In order to compare force generation among different cell lines and stiffness conditions the effect of the varying spreading area needed to be excluded. We found that the total force per cell linearly increased with the spreading area of the cell. Quantitatively this assumption was confirmed by the high linear regression coefficient of R^2^ > 0.45 in all cases (Figure 3A and Supplemental Figure 1A). Therefore, we reasoned that the mean force per pillar, which takes into account the number of deflected pillars depending on the spreading area, resulted in a robust (R^2^ < 0.01) and unbiased measure for cellular force generation (Supplemental Figure 1B). In what follows, traction forces are reported as mean force per pillar under the cell.

**Figure 3.**
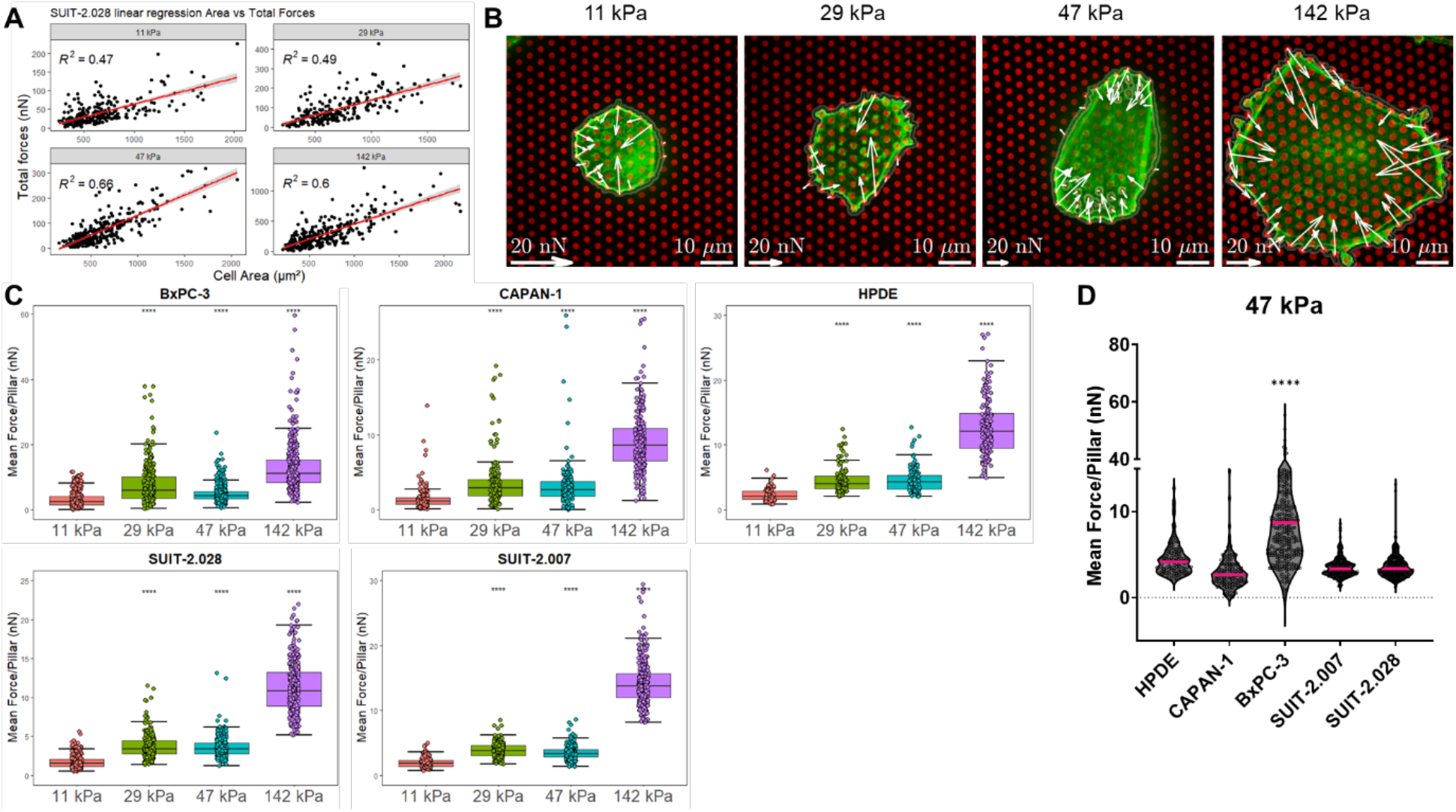
PDAC cell traction forces increase with substrate stiffness, but not with metastatic potential. (**A**) Representative linear regression of total forces (nN) *vs* spreading area (μm^2^) of SUIT-2.028. All linear regression models of the other cell lines are found in Supplemental Figure 1. In all measurements the regression coefficient was R^2^ > 0.45. (**B**) Representative confocal microscopy images of traction forces of BxPC-3 cells growing on fibronectin-coated pillars. White arrows indicate cellular traction forces on the pillars. (**C**) Boxplots of PDAC cell traction forces expressed as mean force per pillar (nN) (25^th^ and 75^th^ percentiles marked, line at median). Statistical significance was calculated using the softest condition (11 kPa) as reference group. (**D**) Mean force per pillar (nN) of PDAC cell lines of different phenotype. Results from other stiffness values (11, 29, 142 kPa) are shown in Supplemental Figure 3. Each dot of plots in (**A**), (**C**) and (**D**) represents the result from one cell.

For all cell lines studied, force generation increased with pillar stiffness (Figures 3B and 3C), corroborating findings on other cell lines in which a likewise stiffness-dependent response was observed [23,25]. For instance, in the BxPC-3 cell line the mean force per pillar increased from 3.2 ± 0.1 nN (mean ± sem) at a substrate stiffness of 11 kPa, to 13.1 ± 0.5 nN at 142 kPa (Figure 3C and Supplemental Table 3).

Next, we assessed whether the cellular phenotype correlates with increased traction forces. It has been described that prostate, breast and lung metastatic cancer cells show an increased single-cell force generation, compared to their respective epithelial-phenotype cells [12]. However, for our panel of PDAC cells we did not observe the same pattern. When grown on substrates of the same stiffness, the non-tumor cell line HPDE and the epithelial cells CAPAN-1 and SUIT-2.028 showed similar traction forces as compared to the metastatic cells SUIT-2.007 (Figure 3D and Supplemental Table 3). Moreover, the epithelial-like cell line BxPC-3 showed a wide fluctuation in forces reaching the highest values compared to all other cell lines investigated.

### Chemoresistant PDAC cells display an altered force generation

Whether PDAC cells with acquired chemoresistance have an altered mechanical characteristic has so far not been investigated. In particular, to date no information is available for PDAC paclitaxel-resistant models. The latter is of interest for mechanobiological studies, as the drug paclitaxel affects microtubule polymerization and cytoskeleton rearrangements. We speculated that the continuous exposure to paclitaxel could alter the mechanobiology of PDAC cells. Therefore, we investigated in three gemcitabine-resistant (GR), and three paclitaxel-resistant (PR) cells [16], whether force generation was affected upon acquired drug-resistance and the concurrent genetic changes that come with resistance. The cell line SUIT-2.028 has an epithelial-like phenotype, while PATU-T and SUIT-2.007 were representative of the mesenchymal phenotype. Given that no significant difference was observed between the intermediate stiffness values (29-47 kPa), and that the stiffest value investigated (142 kPa) has less physiological relevance for PDAC, in what follows we proceeded in comparing the soft and stiff substrates of 11 and 47 kPa only. We first confirmed, as for the non-resistant cells, that PDAC chemoresistant cells had variable cell spreading area, and that the mean force per pillar was a faithful measure of traction forces (see Supplemental Table 4 and Supplemental Figure 4).

Force generation was evaluated on soft and stiff elastic μPAs of 11 kPa and 47 kPa, respectively. For all cell lines we observed an increased force generation for paclitaxel-resistant (PR) *vs* the parental (WT) cells, independent of stiffness (Figure 4 and Supplemental table 5). For the gemcitabine-resistant (GR) cells, both the SUIT-2.028 and SUIT-2.007 cells applied higher traction forces as compared to the parental clones. On the soft substrate (11 kPa) the mean force per pillar was: 1.5 ± 0.1 nN (SUIT-2.028 WT, mean ± sem) *vs* 2.5 ± 0.1 nN (SUIT-2.028 GR) and 2.0 ± 0.1 nN (SUIT-2.007 WT) *vs* 2.2 ± 0.1 nN (SUIT-2.007 GR). On the stiff substrate (47 kPa) the mean force per pillar was: 4.5 ± 0.1 nN (SUIT-2.028 WT) *vs* 5.9 ± 0.1 nN (SUIT-2.028 GR) and 6.3 ± 0.2 nN (SUIT-2.007 WT) vs 8.1 ± 0.3 nN (SUIT-2.007 GR). However, no significant difference was observed for PATU-T GR and WT in both stiffness conditions (Supplemental Table 5). Together, these results indicate that PR-cells apply higher traction forces on both soft and stiff substrates, while in GR-cells an increase in traction force was observed in SUIT-2.028 and SUIT-2.007, but not in PATU-T.

**Figure 4.**
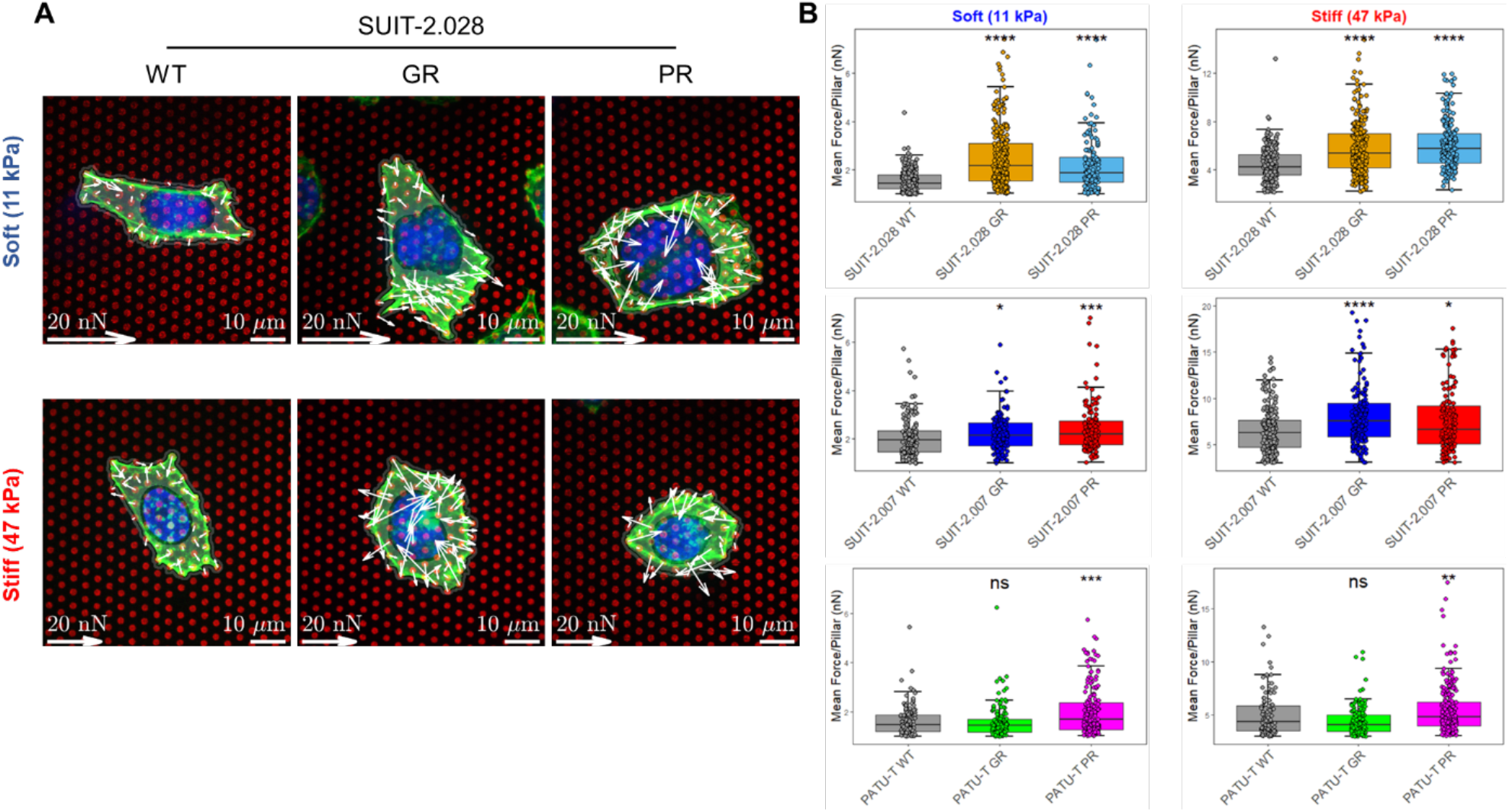
Chemoresistant PDAC cells apply higher traction forces. (**A**) Representative confocal microscopy images of traction forces of SUIT-2.028 WT, GR and PR cells growing on fibronectin-coated soft (11 kPa) and stiff (47 kPa) pillars. White arrows indicate traction forces that cells applied to deflect the pillars. (**B**) Boxplots of PDAC cell traction forces on soft and stiff pillars on the left and right, respectively, expressed as mean force per pillar (nN) (25^th^ and 75^th^ percentiles marked, and line at median). Statistical significance was calculated using the parental cells (WT) as reference group. Each dot represents one cell analyzed.

### Chemoresistant PDAC cells demonstrate distinct migratory behavior compared to their parental cells

Migration and force application are tightly intertwined, as they both rely on the activity of the cytoskeleton [26]. Given the change in force application observed in resistant cells, we wondered whether those would be likewise reflected in the migration and invasive potential of the chemoresistant phenotypes in a single-cell motility assay. Cell nuclei were stained with Hoechst33342 and exposed to 5 µM Verapamil for 1 hour. Verapamil was used to inhibit the efflux of Hoechst, as we earlier found that PR cells overexpress the membrane transporter ABCB1 [16], and thus would quickly lose their Hoechst staining. Before cell plating, substrates were coated by either collagen or fibronectin, as those ECM proteins represent the most abundant proteins in the PDAC microenvironment [27,28]. Stained nuclei were subsequently imaged with confocal fluorescent microscopy at intervals of 12 minutes for 16 hours. Multiple parameters such as single-cell velocity, diffusion constant, diffusive fraction, and directionality were extracted from tracking individual cells over-time (for details see the Supplemental information).

The velocity of chemoresistant PDAC cells was in general different from their parental clones (Figures 5A and 5D). GR-resistant cells were characterized by a slight reduction of their velocity on both substrates. Paclitaxel resistance resulted in a more pronounced, yet also clear cell-specific response. For PATU-T and SUIT-2.028 PR cells the velocity almost halved compared to parental (WT) on both substrates. Conversely, for SUIT-2.007 PR the velocity almost doubled, consistent for both substrates. It appeared that velocity, which characterizes the active, directed part of cell motility is clearly, yet differentially, altered in chemoresistant cells.

**Figure 5.**
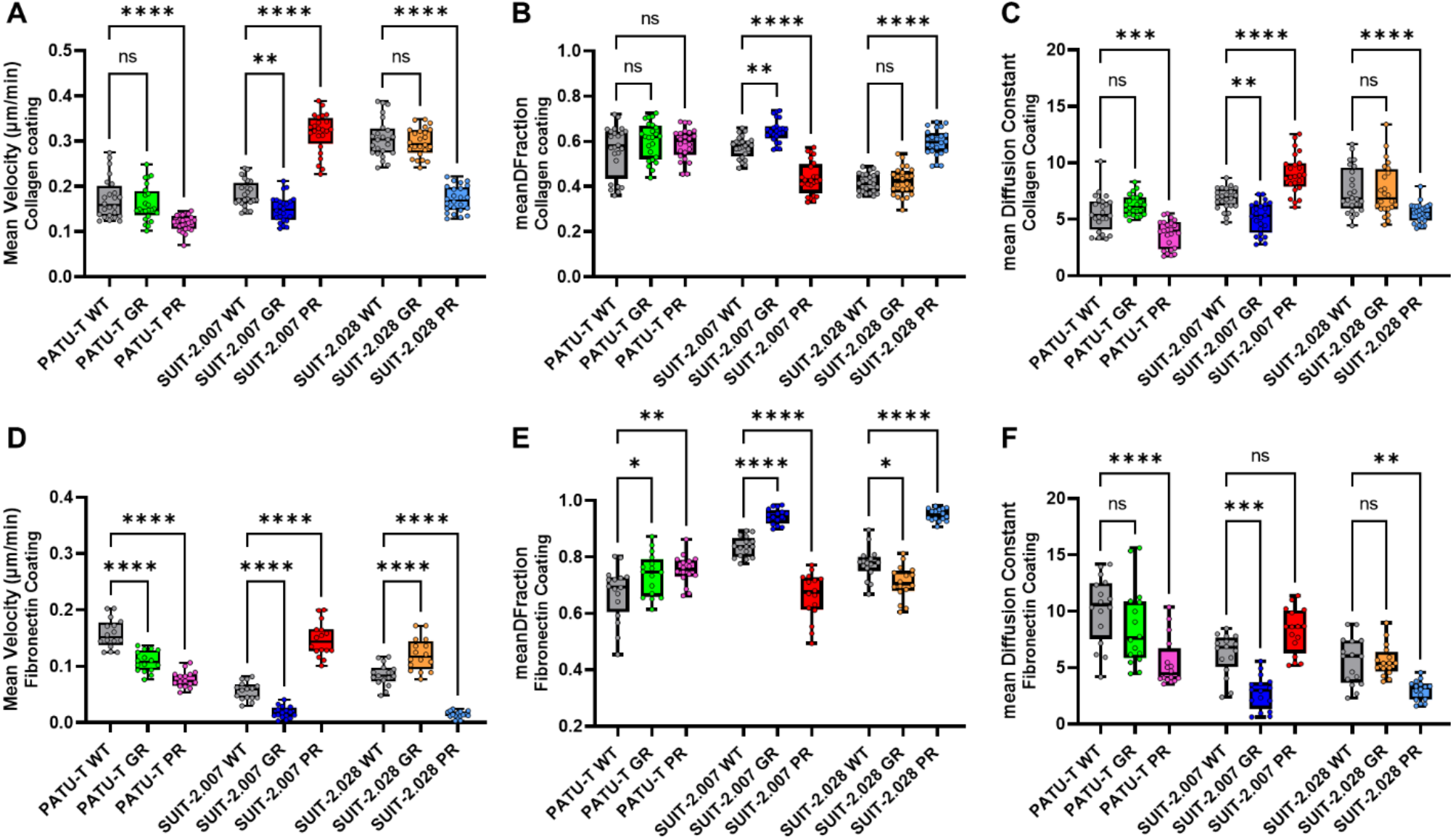
PDAC resistant single-cell migration is different from parental cells. PDAC single-cell mean velocity (μm/min), growing on (**A**) collagen-coated or (**D**) fibronectin-coated wells. Directionality of PDAC cells migration trajectories growing on collagen-coated wells, expressed as (**B**) diffusive fraction = DF and (**C**) diffusion constant. Directionality of PDAC cells migration trajectories growing on fibronectin-coated wells, expressed as (**E**) DF fraction and (**F**) diffusion constant. All conditions are represented as boxplots with 25^th^ and 75^th^ percentiles marked, and line at median. The statistical significance was calculated using the parental cells (WT) as reference group. Each dot represents the population mean for one section of the well.

A measure of the general activity of cells is their diffusional motility characterized by a diffusion constant. In our experiments the diffusion constant followed the pattern of the velocity (Figures 5C and 5F). It should be noted that treatment by 5 µM Verapamil to block the ABCB1 transporter did not affect the migration patterns (Supplemental Figure 5A), while the positive control, bosutinib, which is known to efficiently inhibited migration (Supplemental Figure 5B), resulted in velocity values close to zero.

Given that cell motility was characterized by two modes, active directed motion and diffusion, we further analyzed which of the two modes dominated. For that we defined a diffusive fraction (*DFraction*), which quantifies the fraction of diffusion on the total mean squared-displacement at a timelag of 240 min. *DFraction* approaches unity for purely diffusive motion, and reaches zero for purely directed motion. For all cells and conditions, mobility was diffusion-dominated with *DFraction* > 0.5 (Figure 5 B-E). For most conditions the diffusive fraction only gradually changed, with the pronounced difference for two cell lines resistant to paclitaxel. In detail, where SUIT-2.028 PR cells lost all the active directional part of their motility (*DFraction* = 0.59 ± 0.01), SUIT-2.007 PR cells motility increased the active, directional part of their motility (*DFraction* = 0.44 ± 0.02) (Figures 5B and 5E).

### Migration and force application of PDAC cells in a 3D extracellular matrix

Physiologically, tumor cells move in the three-dimensional tumor-microenvironment (TME). In the TME, cells come into contact with extracellular matrix proteins (collagen, hyaluronic acid, laminin, fibronectin, and others), and with other types of cells (fibroblasts, immune cells, and others). Here we mimicked this complex *in vivo* situation by spheroids of each cell line which were micro-injected in type-1 collagen hydrogels to study the behavior of PDAC cells in a three-dimensional (3D) context resembling the TME. Twenty-four hours after injection, collagen fiber alignment was measured as resemblance of cellular force generation with confocal reflection microscopy. After 48 hours, 3D cell migration was measured as the ratio between the area of the spheroid in z-projection between day-2 and day-0 (Figure 6A). Corroborating the single-cell studies above, resistant cell motility in 3D did not significantly differ from their parental clones, with the exception of SUIT-2.028 GR. Specifically, for the SUIT-2.028 GR cells the relative spheroid area was significantly larger compared to SUIT-2.028 WT (2.5 ± 0.9 *vs*. 1.5 ± 0.9) (Figure 6C and Supplemental Table 6).

**Figure 6.**
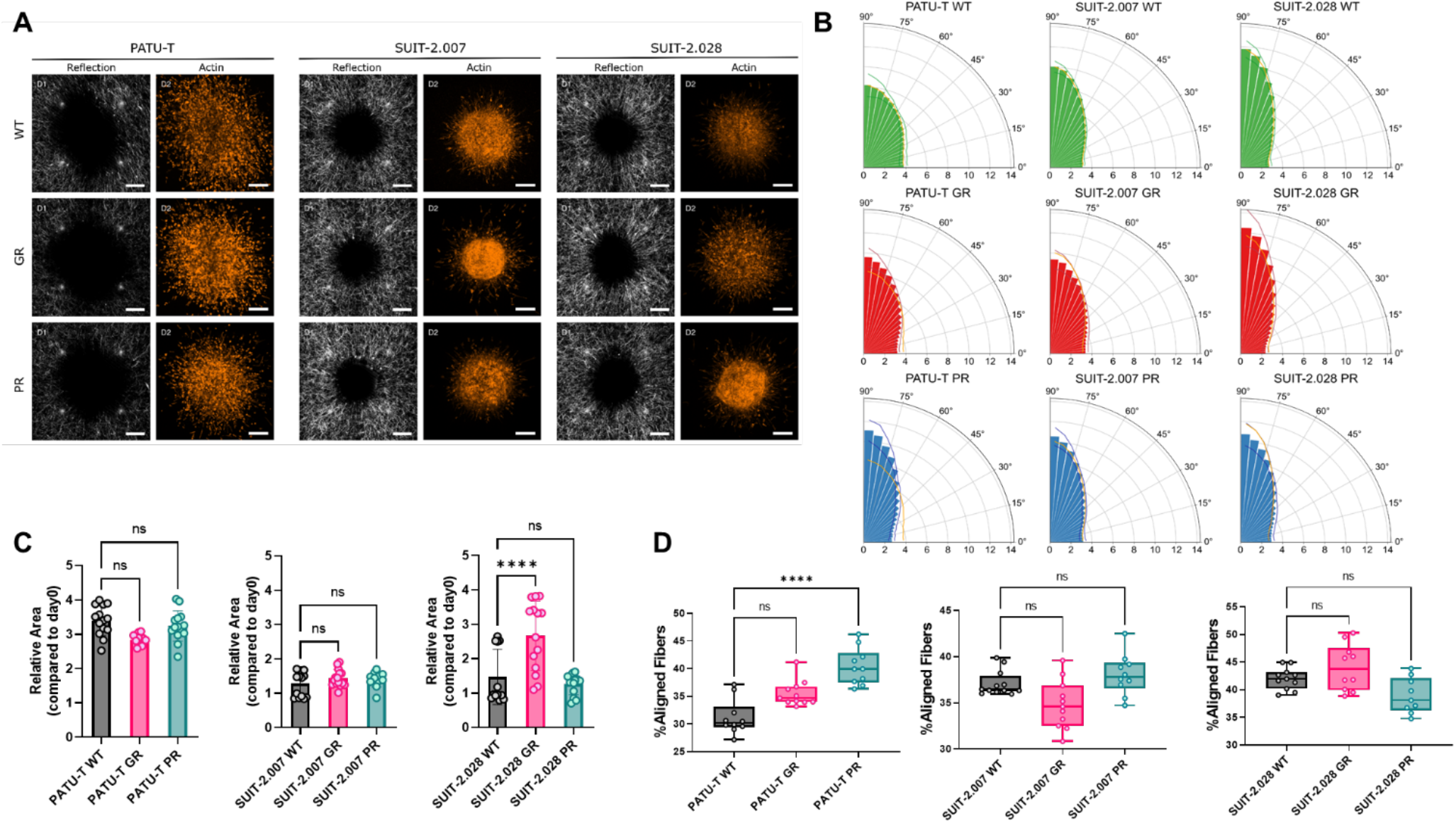
PDAC chemoresistant cell ability for 3D ECM-remodeling and invasion. (**A**) Representative collagen fibers alignment obtained by reflection microscopy (left columns) and actin (right columns) of PDAC chemoresistant spheroids. (**B**) Polar plots representing the percentage of frequency of distribution of collagen fibers angles from 0° to 90°. Each bar represents the % of fibers in a sector of 5 degrees, expressed as the mean of 2 biological replicates, with at least 4 technical replicates. Lines of the darker shade of the bars represent the upper and lower bounds of SD of the % of collagen fibers in each sector. Orange lines represent the mean % of frequency of the respective WT for each cell line, to facilitate the comparison. (**C**) Relative area covered by spheroids after 2 days. Dots represent the value of individual spheroids. (**D**) Percentage of aligned fibers, defined as fibers comprised between the angles of 72.5 and 90, which means that those fibers are perpendicular to the closest point of the spheroid. Dots represent the value of individual spheroids.

The differential force application that we observed in 2D on the uPAs, predicted a differential ability of cells in aligning/remodeling collagen fibers around the spheroids. We therefore measured the angle of the fibers with the closest point to the core of the spheroid. Before micro-injection the collagen fibers were randomly oriented (Supplemental Figure 6). We considered the fibers as aligned for angles between 72.5 and 90 degrees with respect to the tangent of the spheroid outline. Only PATU-T PR caused a significant increase in collagen alignment, despite applying more forces than the WT on pillar for all 3 cell lines (Figures 6B and 6D). We also observed a trend towards increased collagen alignment in SUIT-2.028 GR and PATU-T GR (Figure 6D). However, there was a clear difference in the fraction of aligned fibers between the 3 different parental cell lines, with SUIT-2.028 WT aligning more fibers than the SUIT-2.007 and PATU-T (Figures 6B and 6D).

### YAP nuclear translocation and EMT are not related to increased traction forces

To characterize mechano-response in chemoresistant cells we checked whether differences in traction forces and migration patterns were paralleled by epithelial-to-mesenchymal transition (EMT), and to the occurrence of nuclear yes-associated protein (YAP) translocation. EMT is the classical de-differentiation process epithelial cancer cells undergo, resulting in a more aggressive and invasive phenotype. EMT is characterized by a change in expression of specific genetic markers among which a reduced expression of *E-cadherin* (*E-cad*), and an increase in *N-cadherin* (*N-cad*) and *Vimentin* (*Vim*) expression. Interestingly, when comparing the expression of the abovementioned markers in GR and PR clones *vs* parental cells (WT), a cell-line dependent effect was observed. PATU-T PR displayed an epithelial switch, with reduced expression of both *N-cad* and *Vim*, and a 60-fold increase in *E-cad* expression (Figure 7A). Similarly, SUIT-2.007 GR showed a decreased expression of *Vim*, and a trend towards increased *E-cad* (Figure 7A). However, SUIT-2.028 GR showed a 15-fold increase in *E-cad* expression, while SUIT-2.028 PR an increase in *N-cad* (Figure 7A). Collectively, no consistent pattern was observed for EMT switch among different cell lines, which could explain the increased force generation measured for chemoresistant cells.

**Figure 7.**
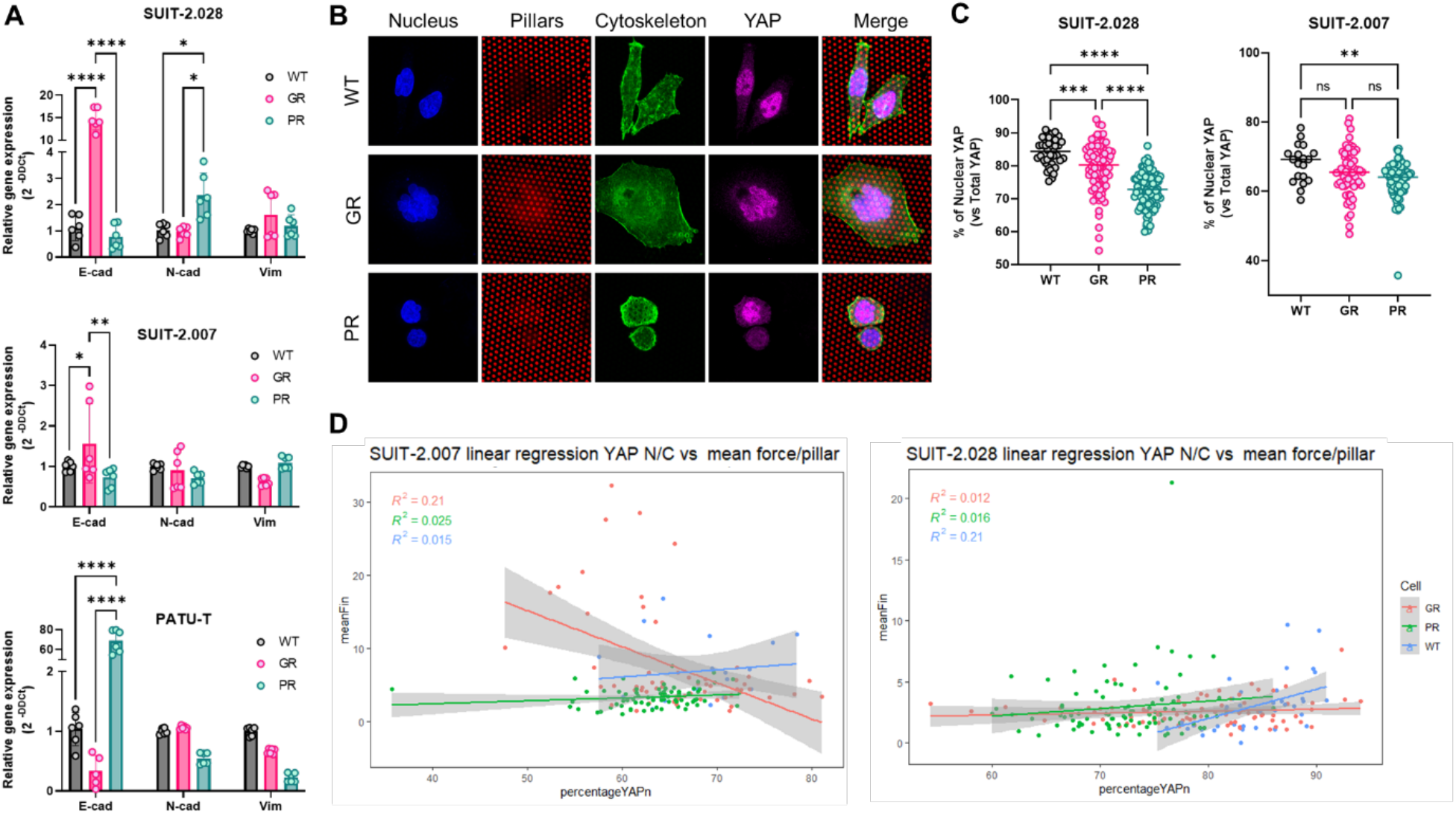
PDAC chemoresistant cell differential mechanobiology do not rely on YAP nuclear translocation nor on EMT switch. (**A**) Relative gene expression of *E-cadherin, N-cadherin* and *Vimentin* as assessed by RT-qPCR. Data are expressed as mean ± sd of three independent experiments. (**B**) Representative confocal microscopy images of SUIT-2.028 cells growing on soft (11 kPa) pillars and stained with YAP. (**C**) YAP nuclear translocation, expressed as % of nuclear YAP over total YAP. (**D**) Linear regression model between the mean force per pillar vs % of nuclear YAP in SUIT-2.007 (left panel) and SUIT-2.028 (right panel). Each dot in (**C**) and (**D**) represents one cell.

Next, we investigated YAP nuclear translocation. This process is triggered upon mechanical stimuli of cells, and marks an active mechanobiology status of the cell [29,30]. Previous studies showed that YAP nuclear translocation is triggered on stiff substrates in mesenchymal stem cells or breast cancer cells [31,32]. Therefore, we analyzed the level of YAP in the nucleus on the softest pillar arrays (11 kPa) to exclude that all YAP was translocated due to stiffness response. Surprisingly, despite both SUIT-2.028/007 GR and PR cells were applying more force than their parental counterparts (WT), we did not find an increase in YAP translocation in the nucleus (Figures 7B and 7C). Conversely, both GR and PR cells showed less YAP in the nucleus compared to WT cells (Figure 7C). To exclude that the increased traction forces are directly related to YAP nuclear translocation we analyzed the correlation between those two quantities in a linear regression model. Interestingly, we found no correlation (R^2^≤0.2) between cellular traction forces and YAP translocation (Figure 7D). Together, our results indicate that the increased traction forces generated by chemoresistant cells do neither rely on EMT nor on YAP signaling.

## Discussion

This is the first study reporting an extensive characterization of mechanobiological features of PDAC chemoresistant cells, indicating that paclitaxel- and gemcitabine-resistant cells apply higher forces, and their motility in 2D and 3D differs from their parental clones.

PDAC acquired chemoresistance is the major cause for poor patient prognosis. Upon drug treatment cancer cells adapt to evade drug-mediated cell death, by modifying signaling pathways, gene expression of drug transporters and other mechanisms [33,34]. However, a better understanding of these biological mechanisms does not fully cover the knowledge of altered cellular processes of PDAC chemoresistance. More precisely, mechanobiology and physical forces play an important role in mediating PDAC chemoresistance. In particular, cells being periodically exposed to cytoskeleton disruptor molecules (e.g. paclitaxel, a microtubule disassembly inhibitor) could acquire differential mechano-transduction patterns. As such, in a previous study of our group we focused on the most widely employed drug for PDAC, i.e. gemcitabine. In this study we reported that PDAC cells acquire resistance to gemcitabine when cultured on stiff substrates, and that cells with acquired chemoresistance have overexpression of the ECM-binding integrin-α2 (ITGA2) [9].

The current study further explored the altered mechanical properties, which we summarize here as mechanobiology, of PDAC cells with acquired resistance to either gemcitabine or paclitaxel, two commonly used drugs in PDAC treatment. Remarkably, we observed that PDAC cells apply more traction forces on stiffer substrates and, upon acquired chemoresistance to either of the drugs, PR or GR showed consistently higher force generation compared to their parental cells. Moreover, chemoresistant cells showed differences in migration, as well as in 3D collagen remodeling and in 3D invasion. However, these altered mechanical and motility features were not reflected by previously reported biological processes, such as an increased YAP nuclear translocation or an EMT switch.

In the current study we used elastic micropillar arrays to measure traction forces generated by a panel of PDAC cell lines. The methodology allowed us to measure cellular contractile forces with a precision below 1 nN [17,35]. When grown on pillars of higher stiffness, PDAC cells displayed a larger spreading area, a finding that is in line with previous studies on fibroblast and endothelial cells using the same experimental settings [18,36]. Additionally, previous studies reported that for fibroblast and other cell types, cell traction forces increase with substrate stiffness [23–25,37]. Here we validated both findings in five PDAC cell lines. Of note, our results were independent on whether cells had an epithelial or mesenchymal phenotype. Furthermore, here we report that PDAC cells with acquired chemoresistance to gemcitabine and paclitaxel adopt an increased contractile behavior as compared to their parental clones. Previous studies mainly focused on investigating PDAC chemoresistance triggered by culturing drug-sensitive cells on substrates with varying stiffness [9,10,38,39]. For instance, Shah and collaborators characterized some mechanical features of PDAC cells with acquired gemcitabine resistance [15], showing a switch to mesenchymal phenotype and an increased migratory/invasive potential. Our results did not validate those findings. This controversy may be explained by variations in experimental conditions. To closely emulate native settings, we indeed cultured cells on ECM-coated substrates and controlled near-native stiffness conditions

EMT and cellular force generation are believed to be closely correlated. For example, metastatic/mesenchymal cancer cells of different tumor types (prostate, breast and lung) showed higher contractile forces [7,12] compared to non-metastatic phenotypes. The correlation between EMT and traction forces in PDAC was first investigated by Nguyen and collaborators, who reported that in PDAC the metastatic potential does not correlate with increased traction forces [5]. The PDAC mesenchymal cell line Hs766 applied less traction forces compared to the epithelial/quasi-mesenchymal PANC-1. Those findings were, in part, corroborated by our results. The epithelial cell line BxPC-3 applied higher forces compared to the mesenchymal SUIT-2.007 cells. However, the non-tumor cells HPDE as well as the epithelial cells (CAPAN-1 and SUIT-2.028) showed no significant differences in traction forces as compared to the mesenchymal cells. This could be, at least in part, explained by the hypothesis proposed by Nguyen and collaborators, suggesting rather than the phenotype, it is the activity of myosin II signaling that orchestrates cellular stiffness and invasion [5].

Prompted by these interesting and contrasting results, we therefore investigated whether cells with acquired chemoresistance had an altered EMT status. We investigated the mesenchymal cell lines PATU-T and SUIT-2.007, as well as the epithelial SUIT-2.028 cells, which despite their origin from a metastatic site, are characterized by a more epithelial phenotype. Regarding traction forces, SUIT-2.007 exhibited the highest forces among all cell types. Yet, PATU-T had lower forces compared to SUIT-2.028. We subsequently investigated whether changes in the expression of EMT genes could explain the observed differences in traction forces between WT and resistant cells. Surprisingly we did not observe a distinct pattern across the cell lines. For some of the cells that showed higher contractile forces, there was a tendency towards an epithelial phenotype switch. Contrarily, a previous study reported a shift towards a mesenchymal status for the gemcitabine-resistant PDAC cells L3.6pl GR, being characterized by a decreased expression of *E-cadherin* and an increase in *Vimentin* [15]. Overall, our findings contradict the preliminary notion that contractile forces and invasive potential correlate with a more mesenchymal phenotype [10,12,14]. This might be explained by the different cellular models employed. In our study we validated results using multiple cell lines with different phenotype (epithelial and mesenchymal), which provides a more robust evidence to our results.

Another robust indicator of activated mechano-transduction in cells, is the translocation of the transcription factor YAP to the nucleus [30]. Previous studies reported that cells growing on stiffer substrates and with an increased invasive potential, showed an elevated YAP nuclear localization [31,32]. Intriguingly, in our study, chemoresistant cells, that applied higher forces, showed less YAP nuclear translocation compared to the parental clones, with no observable correlation with increased contractile forces.

Cell motility is a crucial parameter to characterize the ability of cells to invade tissue during metastatic dissemination. Motility and migration are largely influenced by the surrounding ECM composition, and the rheological properties of tissue [11]. We previously reported that PDAC cells showed increased migration and invasion features, when growing as a monolayer on collagen-coated substrates of controlled stiffness [9]. In the present study, we further explored the role of ECM coating by analyzing single-cell motility on either collagen or fibronectin-coated substrates, the two most abundant ECM proteins found in PDAC. Interestingly, we observed that all chemoresistant cells and their parental clones had the same mobility pattern independently of substrate coating. This finding suggests that PDAC chemoresistant cells underwent some intrinsic (mechano)biological modification, allowing them to migrate differently, rather than adapting to the different ECM substrate. However, when embedded in a 3D matrix, a more controversial behavior was observed: cells appear confined within the collagen matrix and showed a slower invasion rate. We did not find a consistent pattern for the PDAC chemoresistant cell lines. In 2D, the PATU-T cell line exhibited the slowest migration, whereas in 3D the covered area was larger than that of SUIT-2.007 and SUIT-2.028, suggesting that PATU-T cells moved faster through the matrix. One plausible explanation could be attributed to the experimental settings, given the different timescales investigated to measure migration in 2D and 3D environments. The results from spheroids (3D migration) suggest that changes in the covered area were related to the different parental cell line, rather than the drug used to establish resistance.

Regarding the ability to align collagen, gemcitabine resistance led to an increased fiber alignment in SUIT-2.028 GR cells, but a decrease in the SUIT-2.007 GR compared to the WT. Similarly, establishment of paclitaxel resistance resulted in PATU-T PR aligning more fibers, while SUIT-2.028 PR cells exhibited fewer aligned fibers compared to the respective parental cells. It is noteworthy that the GR and PR clones of the same cell line, e.g. SUIT-2.028, had opposite collagen aligning ability when compared to their parental clones. Furthermore, we observed that the ability to align collagen in 3D and the force application on μPAs in 2D do not follow the same trend. Cells applying more forces in 2D are not always able to invade more in 3D, nor are they able to align collagen fibers. This could be in part due to the significant differences in the scaffold stiffness in the two assays. The 2D data have been collected on supports of at least 11 kPa, while collagen hydrogels are typically below 100 Pa [40]. This difference in stiffness could influence the magnitude of force application, as we observed that higher stiffness can trigger an increase in force application. Moreover, it is important to note that the 3D matrices employed are simplistic representations of PDAC tumor microenvironment, which has more ECM proteins than only collagen, thus possibly affecting the differential behavior observed. Not only the stiffness, but the different types of models, such as 2D flat surface versus a 3D matrix with pores, can trigger different types of migration [41], which might not retain the same characteristics. Therefore, the exact mechanism of the controversial behavior in 2D and 3D needs to be elucidated in further studies.

It is important to note that, despite the shared mechanism of resistance development, i.e. the overexpression of ABCB1 transporter [16], force application and migration of PR PDAC cells are not affected in the same way by the prolonged exposure to the drug.

In conclusion, our study characterized multiple mechanical features of PDAC chemoresistant cells and highlighted the importance of including multiple cell models when studying complex physical and biological behaviors. We here reported that to evaluate the effect of drug perturbation on physical parameters tumor heterogeneity plays a crucial role. Therefore, having models that more closely resemble the physiological and physical characteristics of the tumor microenvironment, like tumoroids in a close-to-native TME environment, need to be adopted for a deeper understanding of PDAC mechanobiology.

## References

1. Miller, K.D.; Nogueira, L.; Devasia, T.; Mariotto, A.B.; Yabroff, K.R.; Jemal, A.; Kramer, J.; Siegel, R.L. Cancer Treatment and Survivorship Statistics, 2022. CA. Cancer J. Clin. 2022, 72, 409–436, doi:10.3322/caac.21731.

2. Puik, J.R.; Swijnenburg, R.-J.; Kazemier, G.; Giovannetti, E. Novel Strategies to Address Critical Challenges in Pancreatic Cancer. Cancers 2022, 14, 4115, doi:10.3390/cancers14174115.

3. Garajová, I.; Peroni, M.; Gelsomino, F.; Leonardi, F. A Simple Overview of Pancreatic Cancer Treatment for Clinical Oncologists. Curr. Oncol. 2023, 30, 9587–9601, doi:10.3390/curroncol30110694.

4. Coppola, S.; Carnevale, I.; Danen, E.H.J.; Peters, G.J.; Schmidt, T.; Assaraf, Y.G.; Giovannetti, E. A Mechanopharmacology Approach to Overcome Chemoresistance in Pancreatic Cancer. Drug Resist. Updat. 2017, 31, 43–51, doi:10.1016/j.drup.2017.07.001.

5. Nguyen, A.V.; Trompetto, B.; Tan, X.H.M.; Scott, M.B.; Hu, K.H.; Deeds, E.; Butte, M.J.; Chiou, P.Y.; Rowat, A.C. Differential Contributions of Actin and Myosin to the Physical Phenotypes and Invasion of Pancreatic Cancer Cells. Cell. Mol. Bioeng. 2020, 13, 27–44, doi:10.1007/s12195-019-00603-1.

6. Fan, Y.; Sun, Q.; Li, X.; Feng, J.; Ao, Z.; Li, X.; Wang, J. Substrate Stiffness Modulates the Growth, Phenotype, and Chemoresistance of Ovarian Cancer Cells. Front. Cell Dev. Biol. 2021, 9, 718834, doi:10.3389/fcell.2021.718834.

7. Molter, C.W.; Muszynski, E.F.; Tao, Y.; Trivedi, T.; Clouvel, A.; Ehrlicher, A.J. Prostate Cancer Cells of Increasing Metastatic Potential Exhibit Diverse Contractile Forces, Cell Stiffness, and Motility in a Microenvironment Stiffness-Dependent Manner. Front. Cell Dev. Biol. 2022, 10, 932510, doi:10.3389/fcell.2022.932510.

8. Nabavizadeh, A.; Payen, T.; Iuga, A.C.; Sagalovskiy, I.R.; Desrouilleres, D.; Saharkhiz, N.; Palermo, C.F.; Sastra, S.A.; Oberstein, P.E.; Rosario, V.; et al. Noninvasive Young’s Modulus Visualization of Fibrosis Progression and Delineation of Pancreatic Ductal Adenocarcinoma (PDAC) Tumors Using Harmonic Motion Elastography (HME) in Vivo. Theranostics 2020, 10, 4614–4626, doi:10.7150/thno.37965.

9. Gregori, A.; Bergonzini, C.; Capula, M.; Mantini, G.; Khojasteh-Leylakoohi, F.; Comandatore, A.; Khalili-Tanha, G.; Khooei, A.; Morelli, L.; Avan, A.; et al. Prognostic Significance of Integrin Subunit Alpha 2 (ITGA2) and Role of Mechanical Cues in Resistance to Gemcitabine in Pancreatic Ductal Adenocarcinoma (PDAC). Cancers 2023, 15, 628, doi:10.3390/cancers15030628.

10. Rice, A.J.; Cortes, E.; Lachowski, D.; Cheung, B.C.H.; Karim, S.A.; Morton, J.P.; Del Río Hernández, A. Matrix Stiffness Induces Epithelial–Mesenchymal Transition and Promotes Chemoresistance in Pancreatic Cancer Cells. Oncogenesis 2017, 6, e352–e352, doi:10.1038/oncsis.2017.54.

11. Kumar, S.; Weaver, V.M. Mechanics, Malignancy, and Metastasis: The Force Journey of a Tumor Cell. Cancer Metastasis Rev. 2009, 28, 113–127, doi:10.1007/s10555-008-9173-4.

12. Kraning-Rush, C.M.; Califano, J.P.; Reinhart-King, C.A. Cellular Traction Stresses Increase with Increasing Metastatic Potential. PLoS ONE 2012, 7, e32572, doi:10.1371/journal.pone.0032572.

13. Karamitopoulou, E. Role of Epithelial-Mesenchymal Transition in Pancreatic Ductal Adenocarcinoma: Is Tumor Budding the Missing Link? Front. Oncol. 2013, 3, doi:10.3389/fonc.2013.00221.

14. Zheng, X.; Carstens, J.L.; Kim, J.; Scheible, M.; Kaye, J.; Sugimoto, H.; Wu, C.-C.; LeBleu, V.S.; Kalluri, R. Epithelial-to-Mesenchymal Transition Is Dispensable for Metastasis but Induces Chemoresistance in Pancreatic Cancer. Nature 2015, 527, 525–530, doi:10.1038/nature16064.

15. Shah, A.N.; Summy, J.M.; Zhang, J.; Park, S.I.; Parikh, N.U.; Gallick, G.E. Development and Characterization of Gemcitabine-Resistant Pancreatic Tumor Cells. Ann. Surg. Oncol. 2007, 14, 3629–3637, doi:10.1245/s10434-007-9583-5.

16. Bergonzini, C.; Gregori, A.; Hagens, T.M.S.; Van Der Noord, V.E.; Van De Water, B.; Zweemer, A.J.M.; Coban, B.; Capula, M.; Mantini, G.; Botto, A.; et al. ABCB1 Overexpression through Locus Amplification Represents an Actionable Target to Combat Paclitaxel Resistance in Pancreatic Cancer Cells. J. Exp. Clin. Cancer Res. 2024, 43, 4, doi:10.1186/s13046-023-02879-8.

17. Van Hoorn, H.; Harkes, R.; Spiesz, E.M.; Storm, C.; Van Noort, D.; Ladoux, B.; Schmidt, T. The Nanoscale Architecture of Force-Bearing Focal Adhesions. Nano Lett. 2014, 14, 4257–4262, doi:10.1021/nl5008773.

18. Eckert, J.; Abouleila, Y.; Schmidt, T.; Mashaghi, A. Single Cell Micro-Pillar-Based Characterization of Endothelial and Fibroblast Cell Mechanics. 2021.

19. Schmidt, T.; Schütz, G.J.; Baumgartner, W.; Gruber, H.J.; Schindler, H. Imaging of Single Molecule Diffusion. Proc. Natl. Acad. Sci. 1996, 93, 2926–2929, doi:10.1073/pnas.93.7.2926.

20. Truong, H.H.; De Sonneville, J.; Ghotra, V.P.S.; Xiong, J.; Price, L.; Hogendoorn, P.C.W.; Spaink, H.H.; Van De Water, B.; Danen, E.H.J. Automated Microinjection of Cell-Polymer Suspensions in 3D ECM Scaffolds for High-Throughput Quantitative Cancer Invasion Screens. Biomaterials 2012, 33, 181–188, doi:10.1016/j.biomaterials.2011.09.049.

21. Hou, Y.; Konen, J.; Brat, D.J.; Marcus, A.I.; Cooper, L.A.D. TASI: A Software Tool for Spatial-Temporal Quantification of Tumor Spheroid Dynamics. Sci. Rep. 2018, 8, 7248, doi:10.1038/s41598-018-25337-4.

22. Liu, Y.; Keikhosravi, A.; Mehta, G.S.; Drifka, C.R.; Eliceiri, K.W. Methods for Quantifying Fibrillar Collagen Alignment. In Fibrosis Rittié, L., Ed.; Methods in Molecular Biology; Springer New York: New York, NY, 2017; Vol. 1627, pp. 429–451 ISBN 978-1-4939-7112-1.

23. Califano, J.P.; Reinhart-King, C.A. Substrate Stiffness and Cell Area Predict Cellular Traction Stresses in Single Cells and Cells in Contact. Cell. Mol. Bioeng. 2010, 3, 68–75, doi:10.1007/s12195-010-0102-6.

24. Han, S.J.; Bielawski, K.S.; Ting, L.H.; Rodriguez, M.L.; Sniadecki, N.J. Decoupling Substrate Stiffness, Spread Area, and Micropost Density: A Close Spatial Relationship between Traction Forces and Focal Adhesions. Biophys. J. 2012, 103, 640–648, doi:10.1016/j.bpj.2012.07.023.

25. Balcioglu, H.E.; Harkes, R.; Danen, E.H.J.; Schmidt, T. Substrate Rigidity Modulates Traction Forces and Stoichiometry of Cell–Matrix Adhesions. J. Chem. Phys. 2022, 156, 085101, doi:10.1063/5.0077004.

26. Kashani, A.S.; Packirisamy, M. Cancer Cells Optimize Elasticity for Efficient Migration. R. Soc. Open Sci. 2020, 7, 200747, doi:10.1098/rsos.200747.

27. Liot, S.; Balas, J.; Aubert, A.; Prigent, L.; Mercier-Gouy, P.; Verrier, B.; Bertolino, P.; Hennino, A.; Valcourt, U.; Lambert, E. Stroma Involvement in Pancreatic Ductal Adenocarcinoma: An Overview Focusing on Extracellular Matrix Proteins. Front. Immunol. 2021, 12, 612271, doi:10.3389/fimmu.2021.612271.

28. Tian, C.; Clauser, K.R.; Öhlund, D.; Rickelt, S.; Huang, Y.; Gupta, M.; Mani, D.R.; Carr, S.A.; Tuveson, D.A.; Hynes, R.O. Proteomic Analyses of ECM during Pancreatic Ductal Adenocarcinoma Progression Reveal Different Contributions by Tumor and Stromal Cells. Proc. Natl. Acad. Sci. 2019, 116, 19609–19618, doi:10.1073/pnas.1908626116.

29. Piccolo, S.; Panciera, T.; Contessotto, P.; Cordenonsi, M. YAP/TAZ as Master Regulators in Cancer: Modulation, Function and Therapeutic Approaches. Nat. Cancer 2022, doi:10.1038/s43018-022-00473-z.

30. Dupont, S.; Morsut, L.; Aragona, M.; Enzo, E.; Giulitti, S.; Cordenonsi, M.; Zanconato, F.; Le Digabel, J.; Forcato, M.; Bicciato, S.; et al. Role of YAP/TAZ in Mechanotransduction. Nature 2011, 474, 179–183, doi:10.1038/nature10137.

31. Scott, K.E.; Fraley, S.I.; Rangamani, P. A Spatial Model of YAP/TAZ Signaling Reveals How Stiffness, Dimensionality, and Shape Contribute to Emergent Outcomes. Proc. Natl. Acad. Sci. 2021, 118, e2021571118, doi:10.1073/pnas.2021571118.

32. Caliari, S.R.; Vega, S.L.; Kwon, M.; Soulas, E.M.; Burdick, J.A. Dimensionality and Spreading Influence MSC YAP/TAZ Signaling in Hydrogel Environments. Biomaterials 2016, 103, 314–323, doi:10.1016/j.biomaterials.2016.06.061.

33. Randazzo, O.; Papini, F.; Mantini, G.; Gregori, A.; Parrino, B.; Liu, D.S.K.; Cascioferro, S.; Carbone, D.; Peters, G.J.; Frampton, A.E.; et al. “Open Sesame?”: Biomarker Status of the Human Equilibrative Nucleoside Transporter-1 and Molecular Mechanisms Influencing Its Expression and Activity in the Uptake and Cytotoxicity of Gemcitabine in Pancreatic Cancer. Cancers 2020, 12, 3206, doi:10.3390/cancers12113206.

34. Jain, A.; Bhardwaj, V. Therapeutic Resistance in Pancreatic Ductal Adenocarcinoma: Current Challenges and Future Opportunities. World J. Gastroenterol. 2021, 27, 6527–6550, doi:10.3748/wjg.v27.i39.6527.

35. Li, Z.; Persson, H.; Adolfsson, K.; Abariute, L.; Borgström, M.T.; Hessman, D.; Åström, K.; Oredsson, S.; Prinz, C.N. Cellular Traction Forces: A Useful Parameter in Cancer Research. Nanoscale 2017, 9, 19039–19044, doi:10.1039/C7NR06284B.

36. Balcioglu, H.E.; Van Hoorn, H.; Donato, D.M.; Schmidt, T.; Danen, E.H.J. The Integrin Expression Profile Modulates Orientation and Dynamics of Force Transmission at Cell–Matrix Adhesions. J. Cell Sci. 2015, 128, 1316–1326, doi:10.1242/jcs.156950.

37. Bastounis, E.E.; Yeh, Y.-T.; Theriot, J.A. Subendothelial Stiffness Alters Endothelial Cell Traction Force Generation While Exerting a Minimal Effect on the Transcriptome. Sci. Rep. 2019, 9, 18209, doi:10.1038/s41598-019-54336-2.

38. Xiao, W.; Pahlavanneshan, M.; Eun, C.-Y.; Zhang, X.; DeKalb, C.; Mahgoub, B.; Knaneh-Monem, H.; Shah, S.; Sohrabi, A.; Seidlits, S.K.; et al. Matrix Stiffness Mediates Pancreatic Cancer Chemoresistance through Induction of Exosome Hypersecretion in a Cancer Associated Fibroblasts-Tumor Organoid Biomimetic Model. Matrix Biol. Plus 2022, 14, 100111, doi:10.1016/j.mbplus.2022.100111.

39. Pan, H.; Zhu, S.; Gong, T.; Wu, D.; Zhao, Y.; Yan, J.; Dai, C.; Huang, Y.; Yang, Y.; Guo, Y. Matrix Stiffness Triggers Chemoresistance through Elevated Autophagy in Pancreatic Ductal Adenocarcinoma. Biomater. Sci. 2023, 11, 7358–7372, doi:10.1039/D3BM00598D.

40. Anguiano, M.; Castilla, C.; Maška, M.; Ederra, C.; Peláez, R.; Morales, X.; Muñoz-Arrieta, G.; Mujika, M.; Kozubek, M.; Muñoz-Barrutia, A.; et al. Characterization of Three-Dimensional Cancer Cell Migration in Mixed Collagen-Matrigel Scaffolds Using Microfluidics and Image Analysis. PLOS ONE 2017, 12, e0171417, doi:10.1371/journal.pone.0171417.

41. Friedl, P.; Wolf, K. Plasticity of Cell Migration: A Multiscale Tuning Model. J. Cell Biol. 2010, 188, 11–19, doi:10.1083/jcb.200909003.

