## Supplementary Material for "Altered Mechanobiology of PDAC Cells with Acquired Chemoresistance to Gemcitabine and Paclitaxel"

#### Single-cell motility analysis

The motility of individual PDAC cells was analyzed using a MatLab (MatLab R2018a; MathWorks, Natick, MA, USA) script. For each timepoint images were first thresholded in the respective nuclear marker channel. The center-of-mass positions of all thresholded objects that had the predicted area of a nucleus (10 $\mu m^2$< nuclear area < 400 $\mu m^2$) were determined. From the center-of-mass position data, 2D cell trajectories were constructed using an assignment algorithm described earlier [1]. The mobility of each cell which was observed for at least for 240 min, was further analyzed in terms of the change in the mean-squared displacement (MSD) with lag-time ($t_{lag}$) between two time-points.

We considered two types of movement: one involving diffusion, which is characterized by a diffusion constant D, and a second describing directed active motion characterized by a velocity v [1,2]. In this situation the MSD changes with lag-time were calculated as:

$$MSD(t_{lag}) = 4Dt_{lag} + v^2 t_{lag}^2 \quad (1)$$

In order to characterize the overall motility, we further defined the diffusive fraction $f_D$, as the ratio of the diffusive part of the $MSD_D(t_{lag})$ = 4 D $t_{lag}$, to the total MSD, at a fixed lag-time, $t_D$ = 240 min. The diffusive fraction is given by:

$$f_D = \frac{1}{1+\frac{v^2}{4D} \cdot t_D} \quad (2)$$

**Supplemental Table 1:** list RT-qPCR primers' sequence

| Gene | Forward/Reverse Sequence | Sequence |
|---|---|---|
| **E-cadherin** | Fw | CAATGCCGCCATCGCTTAC |
| | Rv | ATGACTCCTGTGTTCCTGTTAATG |
| **N-cadherin** | Fw | GACAATGCCCCTCAAGTGTT |
| | Rv | CCATTAAGCCGAGTGATGGT |
| **Vimentin** | Fw | GAGAACTTTGCCGTTGAAGC |
| | Rv | GCTTCCTGTAGGTGGCAATC |

**Supplemental Table 2:** spreading area of PDAC cells

| PDAC cells | Stiffness (kPa) | Spreading area (µm² mean ± S.E.M.) |
|---|---|---|
| **HPDE** | 11 | 598 ± 24 |
| | 29 | 582 ± 22 |
| | 47 | 779 ± 28 |
| | 142 | 788 ± 26 |
| **BxPC-3** | 11 | 274 ± 10 |
| | 29 | 295 ± 10 |
| | 47 | 339 ± 11 |
| | 142 | 402 ± 15 |
| **CAPAN-1** | 11 | 221 ± 7 |
| | 29 | 226 ± 8 |
| | 47 | 195 ± 5 |
| | 142 | 476 ± 16 |
| **SUIT-2.028** | 11 | 664 ± 20 |
| | 29 | 704 ± 20 |
| | 47 | 629 ± 18 |
| | 142 | 694 ± 21 |
| **SUIT-2.007** | 11 | 541 ± 19 |
| | 29 | 518 ± 16 |
| | 47 | 694 ± 23 |
| | 142 | 629 ± 18 |

**Supplemental Table 3:** traction forces of PDAC cells

| PDAC cells | Phenotype | Stiffness (kPa) | Number of cells | Traction Force (nN mean ± S.E.M.) |
|---|---|---|---|---|
| **HPDE** | Non-tumor | 11 | 76 | 2,2 ± 0,1 |
| | | 29 | 134 | 4,6 ± 0,2 |
| | | 47 | 144 | 4,5 ± 0,1 |
| | | 142 | 163 | 13,0 ± 0,4 |
| **BxPC-3** | Epithelial | 11 | 357 | 3,2 ± 0,1 |
| | | 29 | 344 | 7,8 ± 0,3 |
| | | 47 | 436 | 5,8 ± 0,2 |
| | | 142 | 307 | 13,1 ± 0,5 |
| **CAPAN-1** | Epithelial | 11 | 27 | 1,4 ± 0,1 |
| | | 29 | 282 | 3,7 ± 0,2 |
| | | 47 | 288 | 3,2 ± 0,2 |
| | | 142 | 253 | 9,0 ± 0,3 |
| **SUIT-2.028** | Epithelial | 11 | 248 | 1,7 ± 0,1 |
| | | 29 | 234 | 3,8 ± 0,1 |
| | | 47 | 293 | 3,6 ± 0,1 |
| | | 142 | 293 | 11,4 ± 0,2 |
| **SUIT-2.007** | Mesenchymal | 11 | 149 | 2,0 ± 0,1 |
| | | 29 | 209 | 3,9 ± 0,1 |
| | | 47 | 228 | 3,5 ± 0,1 |
| | | 142 | 320 | 14,4 ± 0,2 |

**Supplemental Table 4:** spreading area of PDAC chemoresistant cells

| PDAC cells | Chemoresistance status | Stiffness (kPa) | Spreading area ($\mu m^2$ mean ± S.E.M.) |
|---|---|---|---|
| **SUIT-2.028** | WT | 11 | 428 ± 9 |
| | WT | 47 | 480 ± 12 |
| | GR | 11 | 562 ± 18 |
| | GR | 47 | 555 ± 18 |
| | PR | 11 | 372 ± 18 |
| | PR | 47 | 405 ± 16 |
| **SUIT-2.007** | WT | 11 | 522 ± 19 |
| | WT | 47 | 500 ± 15 |
| | GR | 11 | 418 ± 22 |
| | GR | 47 | 371 ± 14 |
| | PR | 11 | 485 ± 24 |
| | PR | 47 | 482 ± 18 |
| **PATU-T** | WT | 11 | 520 ± 18 |
| | WT | 47 | 518 ± 25 |
| | GR | 11 | 621 ± 21 |
| | GR | 47 | 641 ± 28 |
| | PR | 11 | 556 ± 20 |
| | PR | 47 | 498 ± 14 |

**Supplemental Table 5:** traction forces of PDAC chemoresistant cells

| PDAC cells | Chemoresistance status | Stiffness (kPa) | Number of cells | Traction Force (nN mean ± S.E.M.) |
|---|---|---|---|---|
| **SUIT-2.028** | WT | 11 | 280 | 1,5 ± 0,1 |
| | GR | 11 | 289 | 2,5 ± 0,1 |
| | PR | 11 | 152 | 2,1 ± 0,1 |
| | WT | 47 | 248 | 4,5 ± 0,1 |
| | GR | 47 | 284 | 5,9 ± 0,1 |
| | PR | 47 | 222 | 5,9 ± 0,2 |
| **SUIT-2.007** | WT | 11 | 158 | 2,0 ± 0,1 |
| | GR | 11 | 147 | 2,2 ± 0,1 |
| | PR | 11 | 129 | 2,4 ± 0,1 |
| | WT | 47 | 218 | 6,3 ± 0,2 |
| | GR | 47 | 158 | 8,1 ± 0,3 |
| | PR | 47 | 153 | 7,7 ± 0,3 |
| **PATU-T** | WT | 11 | 167 | 1,5 ± 0,1 |
| | GR | 11 | 243 | 1,3 ± 0,1 |
| | PR | 11 | 212 | 1,9 ± 0,1 |
| | WT | 47 | 165 | 4,3 ± 0,2 |
| | GR | 47 | 221 | 3,9 ± 0,1 |
| | PR | 47 | 253 | 5,3 ± 0,2 |

**Supplemental Table 6:** Mean relative area of PDAC spheroids migrating in collagen matrix.

| PDAC cells | Chemoresistance Status | Relative Area (Mean ± S.E.M.) |
|---|---|---|
| **PATU-T** | WT | 3,4 ± 0,3 |
| | GR | 2,8 ± 0,1 |
| | PR | 3,2 ± 0,3 |
| **SUIT-2.007** | WT | 1,3 ± 0,3 |
| | GR | 1,4 ± 0,2 |
| | PR | 1,4 ± 0,2 |
| **SUIT-2.028** | WT | 1,5 ± 0,6 |
| | GR | 2,7 ± 0,7 |
| | PR | 1,3 ± 0,2 |

### Supplemental Figure 1

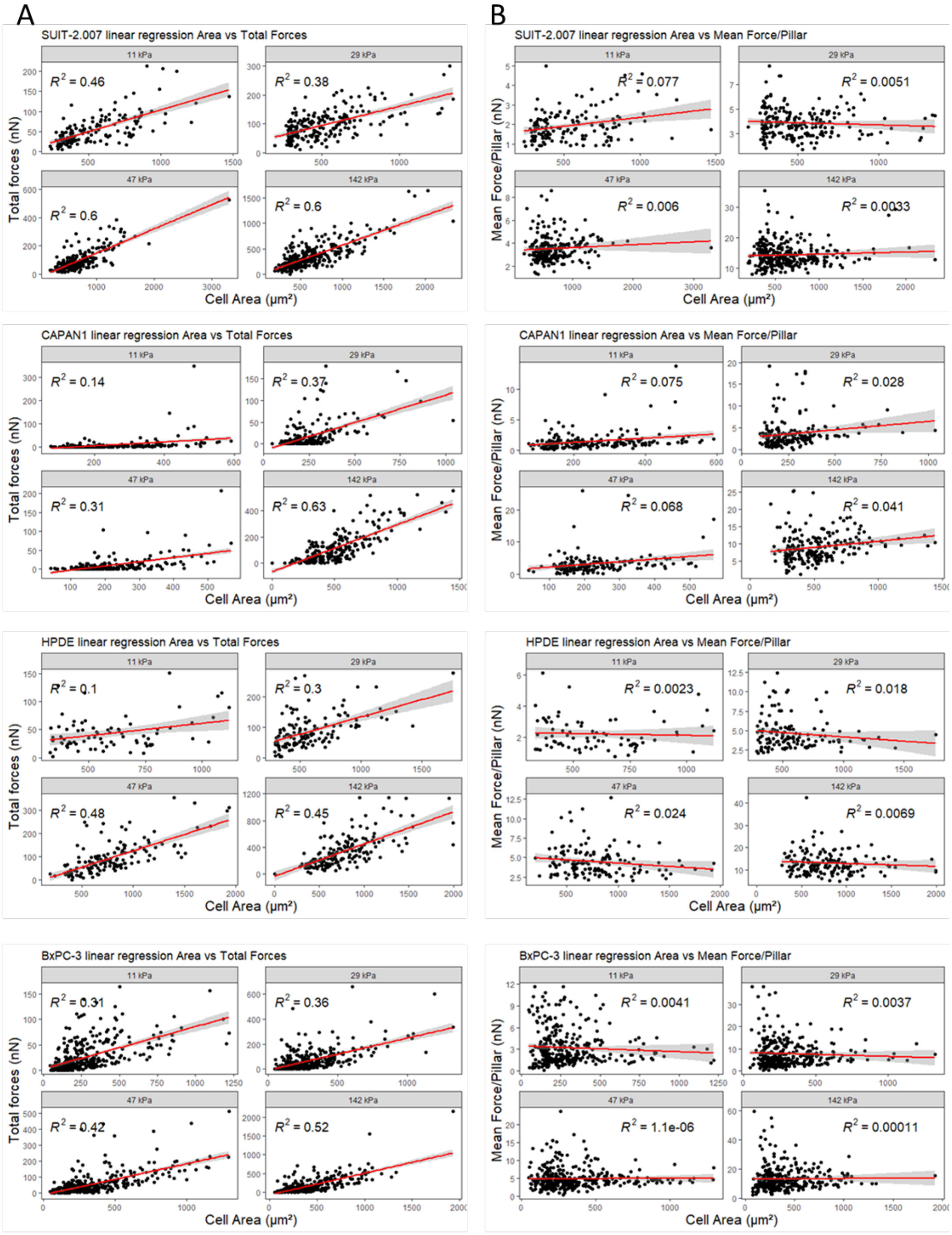


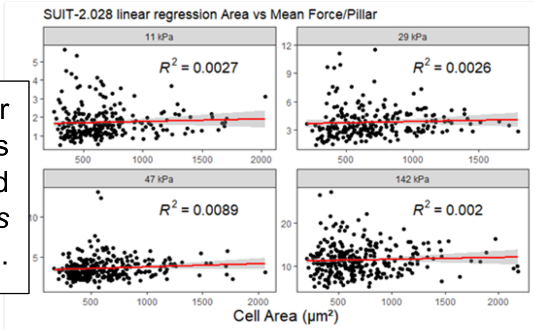


**Supplemental Figure 1:** Linear regression model of (**A**) total forces (nN) *vs* spreading area (µm$^2$), and (**B**) mean force per pillar (nN) *vs* spreading area (µm$^2$) of PDAC cells.

#### Supplemental Figure 2

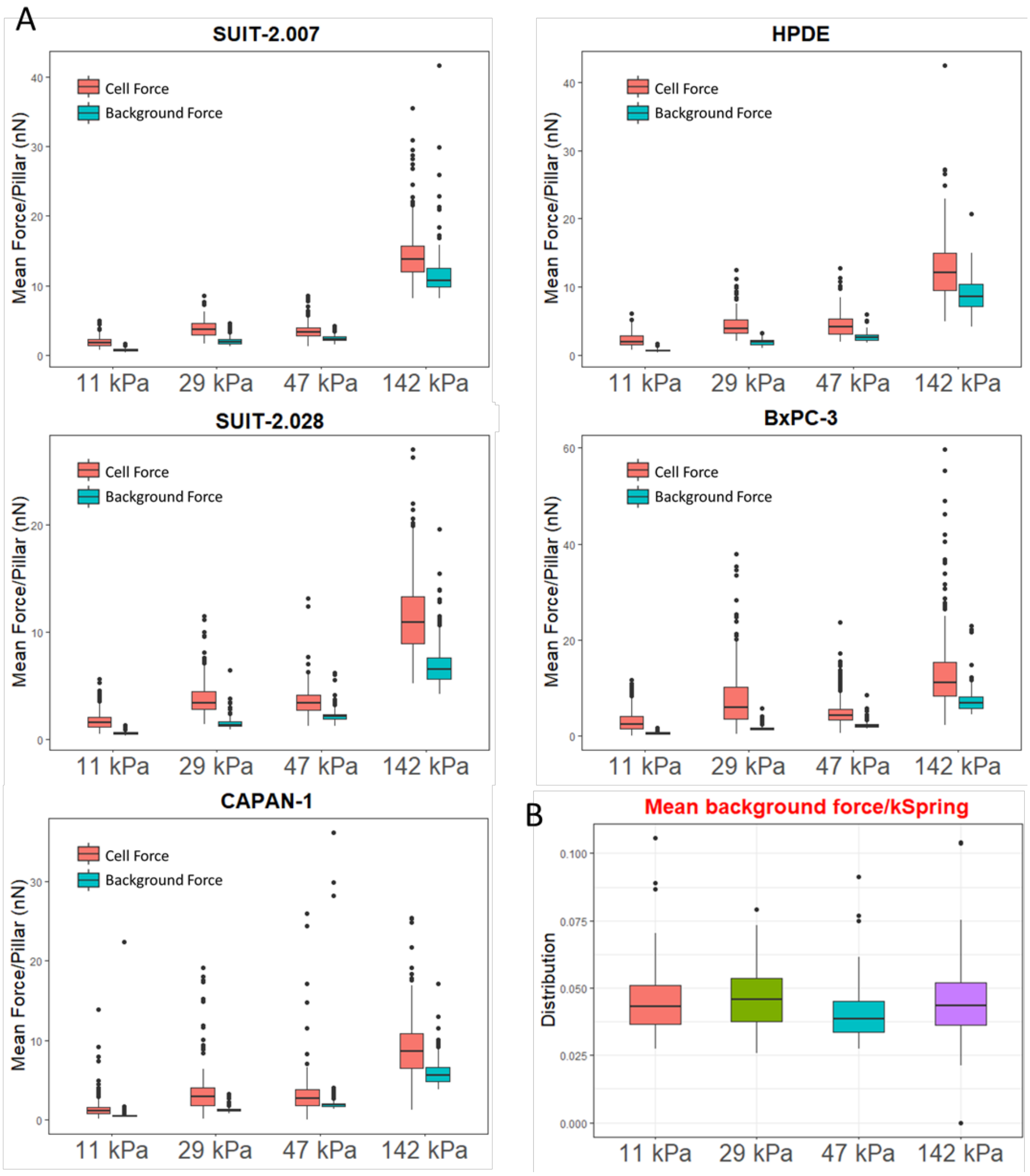


**Supplemental Figure 2:** Pillar ‘background forces’. (**A**) Mean force per pillar (nN) calculated on pillars deflected either under the cell area (orange = cellular force) or outside the cell area (cerulean = ‘background force’). The ‘background force’ is given by the accuracy, at which the center-of-mass of each pillar is determined. It’s value is given by the ratio of the pillar diameter (2 µm), and the square-root of the integrated signal for each pillar [3]. In our experiment the integrated signal was ~2000 cnts, which results in an accuracy of pillar detection of ~50 nm. Multiplication with the respective spring constant results in an apparent background-force. Since we report on force magnitude only, the background-force does not vanish but is finite. In all cases, the cellular forces clearly exceed the background. (**B**) When background-forces are divided by the respective spring constants, the resulting displacements had indeed identical distributions of mean of 0.04 ± 0.01 um (mean ± sd), as predicted from the theory [3].

#### Supplemental Figure 3

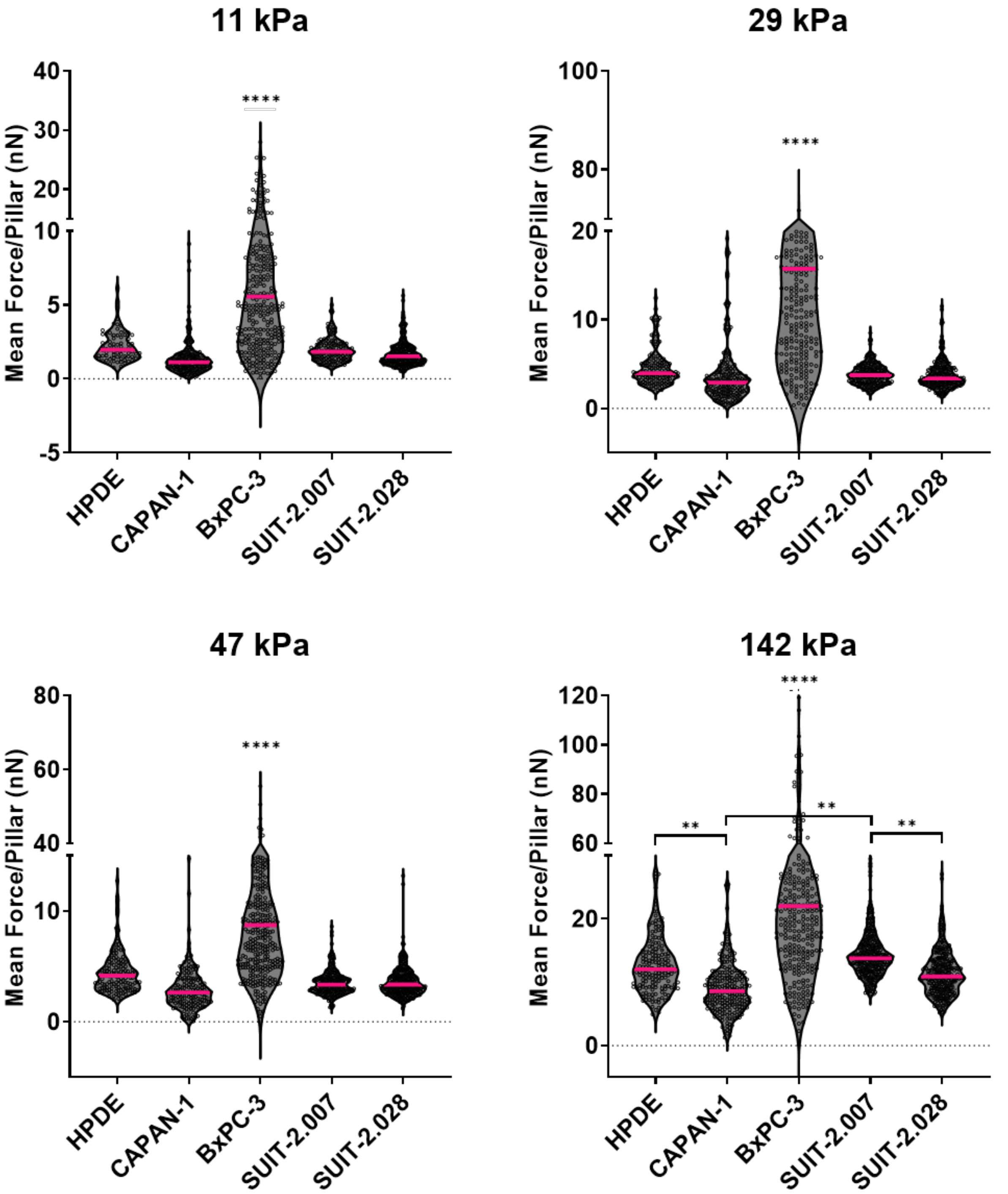


**Supplemental Figure 3:** Traction force of PDAC cells. Mean force per pillar (nN) of different PDAC cell lines growing on pillars with varying stiffness.

#### Supplemental Figure 4

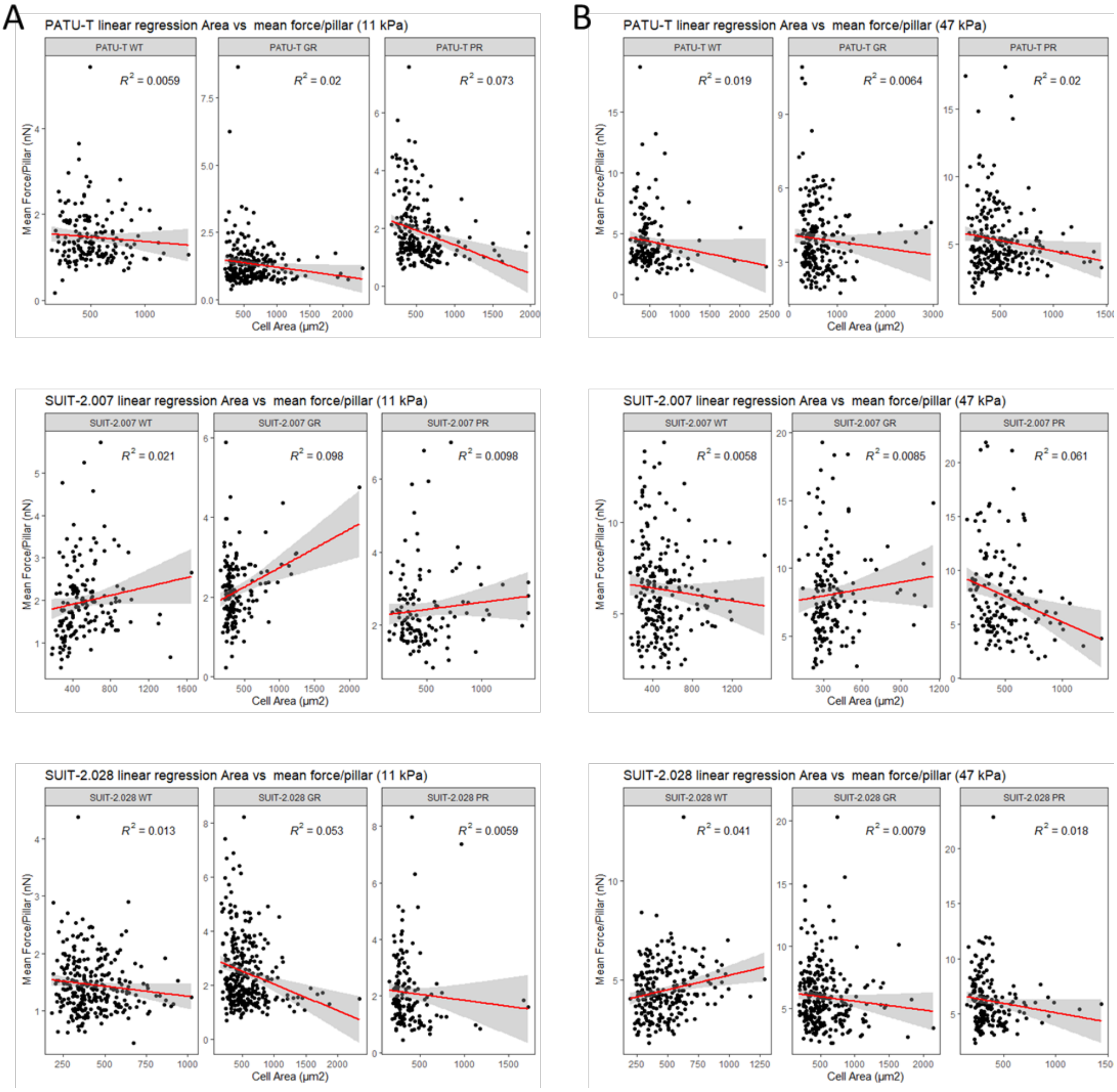


**Supplemental Figure 4:** Linear regression model of the mean force per pillar (nN) *vs* spreading area (µm$^2$) of PDAC chemoresistant cells grown on (**A**) soft (11 kPa), and (**B**) stiff (47 kPa) pillars. Note that in all cases $R^2 \leq 0,02$, indicating that also for PDAC chemoresistant cells the mean force per pillar is uncorrelated to spreading area.

#### Supplemental Figure 5

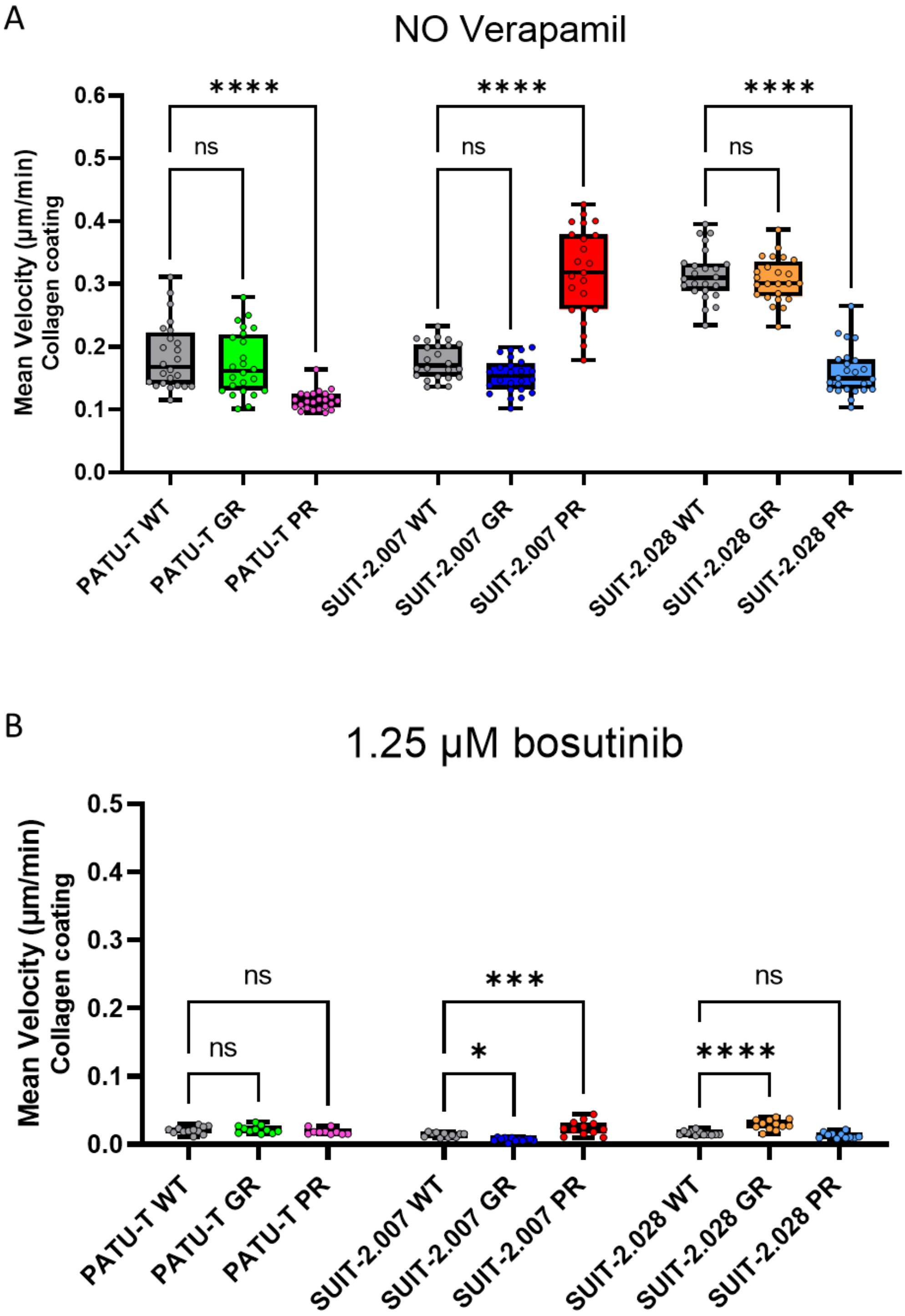


**Supplemental figure 5:** PDAC cell migration is effectively inhibited by the motility-inhibitor bosutinib but not affected by the ABCB1-blocker verapamil. PDAC cell velocity, expressed as mean velocity (µm/min), growing on collagen-coated wells (**A**) untreated (no verapamil), and (**B**) treated with bosutinib (positive control).

#### Supplemental Figure 6

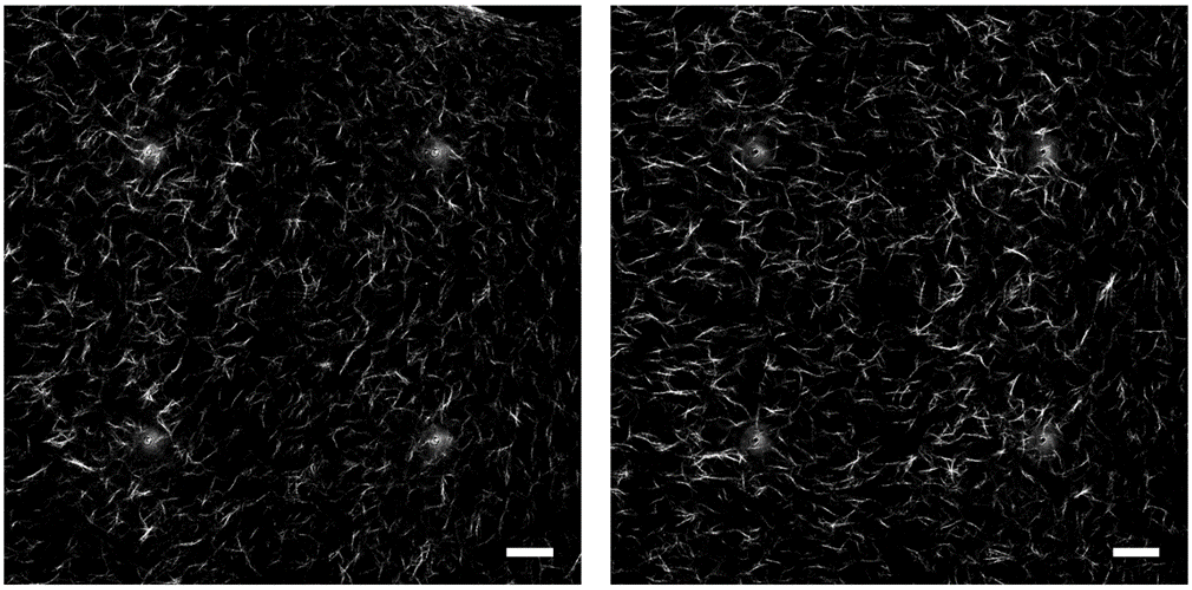

**Supplemental figure 6:** Representative confocal reflection images of single z-planes of empty collagen gels, showing the random orientation of collagen fibers. Scale bar is 200 μm.

#### References


1. Schmidt, T.; Schütz, G.J.; Baumgartner, W.; Gruber, H.J.; Schindler, H. Imaging of Single Molecule Diffusion. *Proc. Natl. Acad. Sci.* **1996**, *93*, 2926–2929, doi:10.1073/pnas.93.7.2926.

2. Kusumi, A.; Sako, Y.; Yamamoto, M. Confined Lateral Diffusion of Membrane Receptors as Studied by Single Particle Tracking (Nanovid Microscopy). Effects of Calcium-Induced Differentiation in Cultured Epithelial Cells. *Biophys. J.* **1993**, *65*, 2021–2040, doi:10.1016/S0006-3495(93)81253-0.

3. Bobroff, N. Position Measurement with a Resolution and Noise-Limited Instrument. *Rev. Sci. Instrum.* **1986**, *57*, 1152–1157, doi:10.1063/1.1138619.
